# Morphological stasis masks ecologically divergent coral species on tropical reefs

**DOI:** 10.1101/2020.09.04.260208

**Authors:** P Bongaerts, IR Cooke, H Ying, D Wels, S Haan den, A Hernandez-Agreda, CA Brunner, S Dove, N Englebert, G Eyal, S Forêt, M Grinblat, KB Hay, S Harii, DC Hayward, Y Lin, M Mihaljević, A Moya, P Muir, F Sinniger, P Smallhorn-West, G Torda, MA Ragan, MJH van Oppen, O Hoegh-Guldberg

**Affiliations:** California Academy of Sciences, San Francisco, California, USA; Global Change Institute, The University of Queensland, Brisbane, Australia; ARC Centre of Excellence for Coral Reef Studies, School of Biological Sciences, The University of Queensland, Brisbane, Australia; Department of Molecular and Cell Biology, James Cook University, Townsville, Queensland, Australia; Centre for Tropical Bioinformatics and Molecular Biology, James Cook University, Townsville, Queensland, Australia; Division of Ecology and Evolution, Research School of Biology, Australian National University, Canberra, Australia; Department of Environmental Sciences and Policy, Central European University, Budapest, Hungary; ARC Centre of Excellence for Coral Reef Studies, James Cook University, Townsville, Australia; Australian Institute of Marine Science, Townsville, Queensland, Australia; The Mina & Everard Goodman Faculty of Life Sciences, Bar-Ilan University, Ramat Gan, Israel; Tropical Biosphere Research Center, University of the Ryukyus, Motobu, Okinawa, Japan; Research School of Computer Science, Australian National University, Canberra, Australia; Science Lab UZH, University of Zurich, Zurich, Switzerland; Department of Biology, University of Konstanz, Konstanz, Germany; Queensland Museum, Townsville, Australia; Institute for Molecular Bioscience, The University of Queensland, Brisbane, Australia; School of BioSciences, the University of Melbourne, Victoria, Australia

## Abstract

Coral reefs are the epitome of species diversity, yet the number of described scleractinian coral species, the framework-builders of coral reefs, remains moderate by comparison. DNA sequencing studies are rapidly challenging this notion by exposing a wealth of undescribed diversity, but the evolutionary and ecological significance of this diversity remains largely unclear. Here, we present an annotated genome for one of the most ubiquitous corals in the Indo-Pacific (*Pachyseris speciosa*), and uncover through a comprehensive genomic and phenotypic assessment that it comprises morphologically indistinguishable, but ecologically divergent cryptic lineages. Demographic modelling based on whole-genome resequencing disproved that morphological crypsis was due to recent divergence, and instead indicated ancient morphological stasis. Although the lineages occur sympatrically across shallow and mesophotic habitats, extensive genotyping using a rapid diagnostic assay revealed differentiation of their ecological distributions. Leveraging “common garden” conditions facilitated by the overlapping distributions, we assessed physiological and quantitative skeletal traits and demonstrated concurrent phenotypic differentiation. Lastly, spawning observations of genotyped colonies highlighted the potential role of temporal reproductive isolation in the limited admixture, with consistent genomic signatures in genes related to morphogenesis and reproduction. Overall, our findings demonstrate how ecologically and phenotypically divergent coral species can evolve despite morphological stasis, and provide new leads into the potential mechanisms facilitating such divergence in sympatry. More broadly, they indicate that our current taxonomic framework for reef-building corals may be scratching the surface of the ecologically relevant diversity on coral reefs, consequently limiting our ability to protect or restore this diversity effectively.

## INTRODUCTION

Tropical coral reefs are known for their high levels of biodiversity, harboring hundreds of thousands of macroscopic and an unknown number of microscopic species (1). Interestingly, the contribution of reef-building corals remains surprisingly moderate with only ~750-850 valid species accepted worldwide (2–5). Under the current taxonomic framework, species are distinguished primarily based on diagnostic skeletal characteristics, an approach that originated from the early days of coral reef science when underwater observations were extremely challenging (6, 7). Skeletal traits at both the corallite and colony level are often highly variable within species and across environments, posing a major challenge to coral taxonomy (8). Conversely, skeletal traits are known to converge and can even obscure deep phylogenetic divergences at the family level (9, 10), highlighting further challenges to the use of the skeletal morphospace as a taxonomic framework. Molecular approaches have helped resolve some of these difficulties by differentiating morphological plasticity from actual species traits (11, 12), confirming species separation in the context of subtle morphological differences (13–15), and clarifying deeper evolutionary relationships within the order (9, 10). However, molecular studies have also uncovered a wealth of undescribed diversity within taxonomic species that cannot be readily explained (11, 16–24).

The notion that scleractinian coral diversity may be far greater than acknowledged through conventional taxonomy is not novel. A seminal review published over two decades ago highlighted the ubiquitous presence of sibling species in the marine realm, and stressed the importance of exploring its ecological relevance (25). Since then, molecular studies have indeed exposed “taxonomically cryptic diversity” (i.e. genetically distinct taxa that have been erroneously classified under a single species name) within many coral genera (21, 26). However, much of this cryptic diversity is being exposed through population genetic studies designed to relate genetic patterns to geography (26), and consequently, it usually remains unclear to what extent they represent phenotypically and ecologically distinct entities [but see (27–31)], including whether lineages are truly morphologically cryptic. In fact, well-studied examples of coral species complexes are characterized by substantial morphological differentiation (14,16,17,20,27), and we still know little about the potential for ecological or phenotypic differentiation when gross morphology is constrained. The traditional use of only a few genetic loci has also impeded an assessment of the evolutionary context of cryptic diversification, and the respective roles of neutral versus selective processes on ecological or reproductive traits (31, 32). Thus, the ecological relevance of cryptic diversity in corals remains poorly understood and is therefore rarely considered in ecological studies or conservation planning.

To address this knowledge gap, we conducted a comprehensive assessment to evaluate the nature of cryptic diversity in *Pachyseris speciosa* (“Serpent Coral”; Figure 1a), one of the most ubiquitous and abundant species across the Indo-Pacific (3). This zooxanthellate coral has one of the widest bathymetric distributions on tropical coral reefs –from close to the surface to lower mesophotic depths (~5-95 m) (33)– and has therefore been the focus of studies assessing its ecological opportunism (34–37). Our initial assessment of its genetic structure indicated the presence of undescribed sympatric lineages, and we therefore used this as an opportunity to explore the extent to which cryptic diversity can obscure genomic, ecological and phenotypic divergence, and how reproductive isolation may be maintained in such closely-related lineages. Specifically, our study (summarized in Figure 1) involved the generation of an annotated genome for *P. speciosa* using long-read sequencing, which we used as a reference for reduced-representation sequencing (nextRAD) of populations ranging from across the geographic and bathymetric distribution of this species. Subsequent whole-genome resequencing (WGS) of representative coral colonies was implemented for demographic modelling, and for the development of a cleaved amplified polymorphic sequence (CAPS) assay to assess ecological, phenotypic and reproductive differentiation in the uncovered lineages from Australia. Overall, our findings demonstrate the potential for ecological and phenotypic divergence under morphological stasis, and highlight that conventional skeleton-based taxonomy may be substantially underestimating the ecologically-relevant diversity of scleractinian corals on tropical coral reefs.

**Figure 1.**
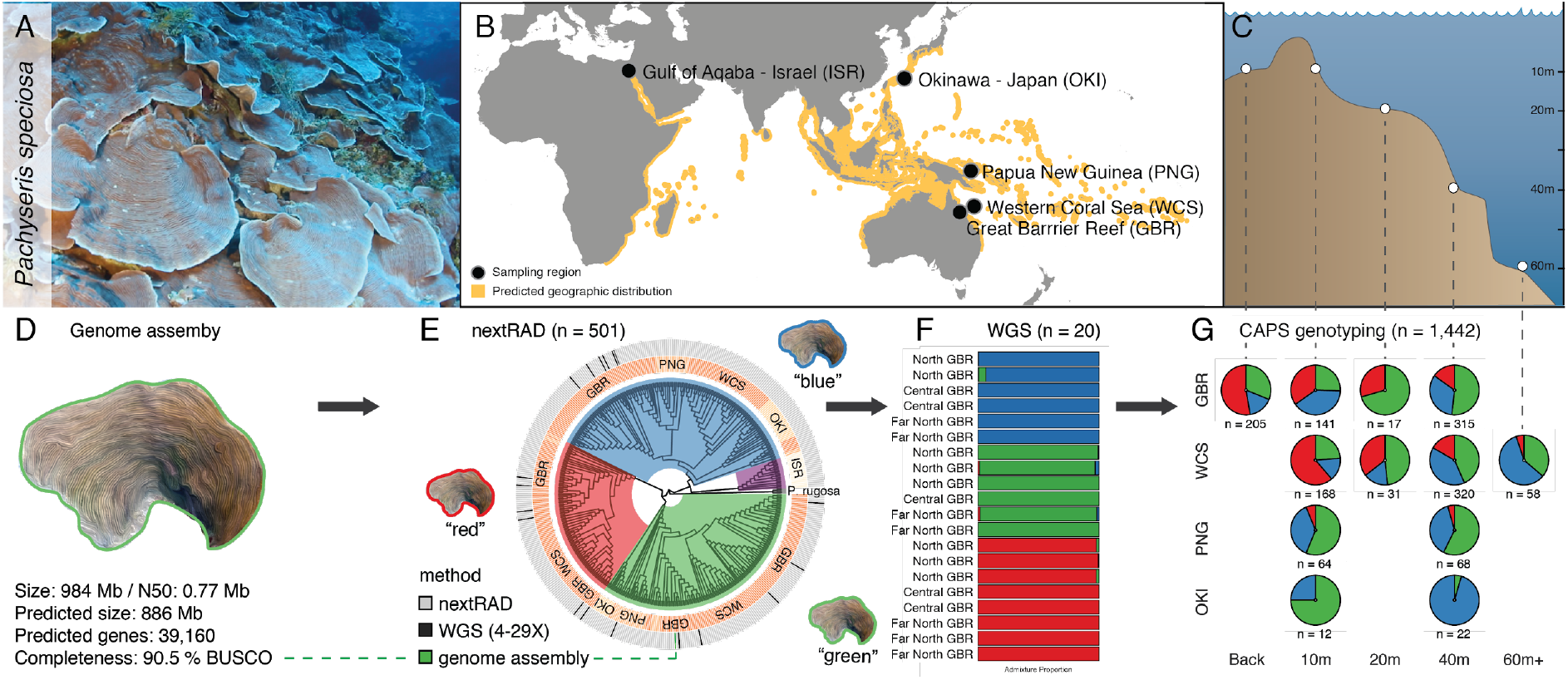
Overview of the study system and sequencing/genotyping approach. (A) *Pachyseris speciosa* at 40 m depth at Holmes Reef (Western Coral Sea). (B) Predicted geographic distribution of *P. speciosa* (*sensu lato*) based on the IUCN Red List (2009), and the location of the five regions included in this study. (C) Sampled habitats and their corresponding water depths (± 3 m). (D) Genome assembly summary statistics. (E) Circular phylogenetic tree summarizing the reduced-representation sequencing (nextRAD) data, and showing the partitioning into four major lineages. Sample region is indicated in the inner surrounding circle, and selection for whole-genome sequencing indicated in the outer surrounding circle. (F) Admixture proportion (PCAngsd) based on whole-genome resequencing (WGS) of representative samples belonging to the three lineages occurring on the Great Barrier Reef. (G) Geographic and ecological distribution of the lineages across habitats and regions (excluding Israel). Assignments are based on the CAPS genotyping assay merged with the nextRAD data (note that assignments for Okinawa are based exclusively on nextRAD data).

## Results and Discussion

### Genome assembly and annotation

We generated a highly contiguous reference genome for *Pachyseris speciosa*, assembled through PacBio single-molecule long-read sequencing (Figure 1d; Table S2). The assembled genome size is 984 Mb, representing one of the largest coral genomes reported to date, and comprising 2,368 contigs with a N50 size of 766.6 kb (largest contig 4.6 Mb; Table S3,4). In total, 39,160 protein-coding genes were predicted – a number comparable to that of two other recently sequenced corals from the “robust” clade [i.e. one of the two major clades of extant corals; (38)] – with the gene model being 90.5% complete and 5.1% partial (based on conserved single-copy metazoan orthologous genes) (Table S5,6,7). *De novo* repeat annotation revealed that 52.2% of the assembly is occupied by repetitive elements, which are generally better-resolved in long-read assemblies (39). The dominance of transposable elements compared with other robust corals indicates that the larger genome size may be substantially driven by expansion of those transposons (38) (Table S8).

### *Pachyseris speciosa* represents a sympatric species complex

The annotated genome assembly was used as a reference for reduced-representation sequencing (nextRAD), where we sequenced *P. speciosa* colonies (n = 501) from >30 sites from Australasia [the Great Barrier Reef (GBR), the Western Coral Sea (WCS), and Papua New Guinea (PNG)], Okinawa (OKI), and Israel (ISR) (Figure 1E, S1; Table S1). Genetic structuring based on principal component and neighbor-joining analyses revealed the presence of four highly divergent lineages (Figures 2, S2). One lineage represents the geographically separated Israel population (Gulf of Aqaba) and indicates the existence of a distinct species in the Red Sea region, corroborating a pattern observed in other corals [e.g. *Stylophora pistillata*; (24, 40–41)]. The other three lineages occurred sympatrically on all of the sampled reefs in Australasia, hereafter arbitrarily referred to as “green”, “blue” and “red” lineage, with the assembled reference genome representing a genotype belonging to the “green” lineage. Subsequent structure analyses using a maximum likelihood framework (snapclust) indicated an optimum between four and six clusters, and the additional two clusters (for k = 5 and 6) correspond to two populations from Okinawa that grouped with respectively the “blue” and “green” lineages but were substantially differentiated from those in eastern Australia (Figures S2,S3). Beyond this expected geographic differentiation of sub-tropical Okinawa, the overall structuring demonstrates the presence of genetically distinct lineages within the currently acknowledged species, *P. speciosa*.

**Figure 2.**
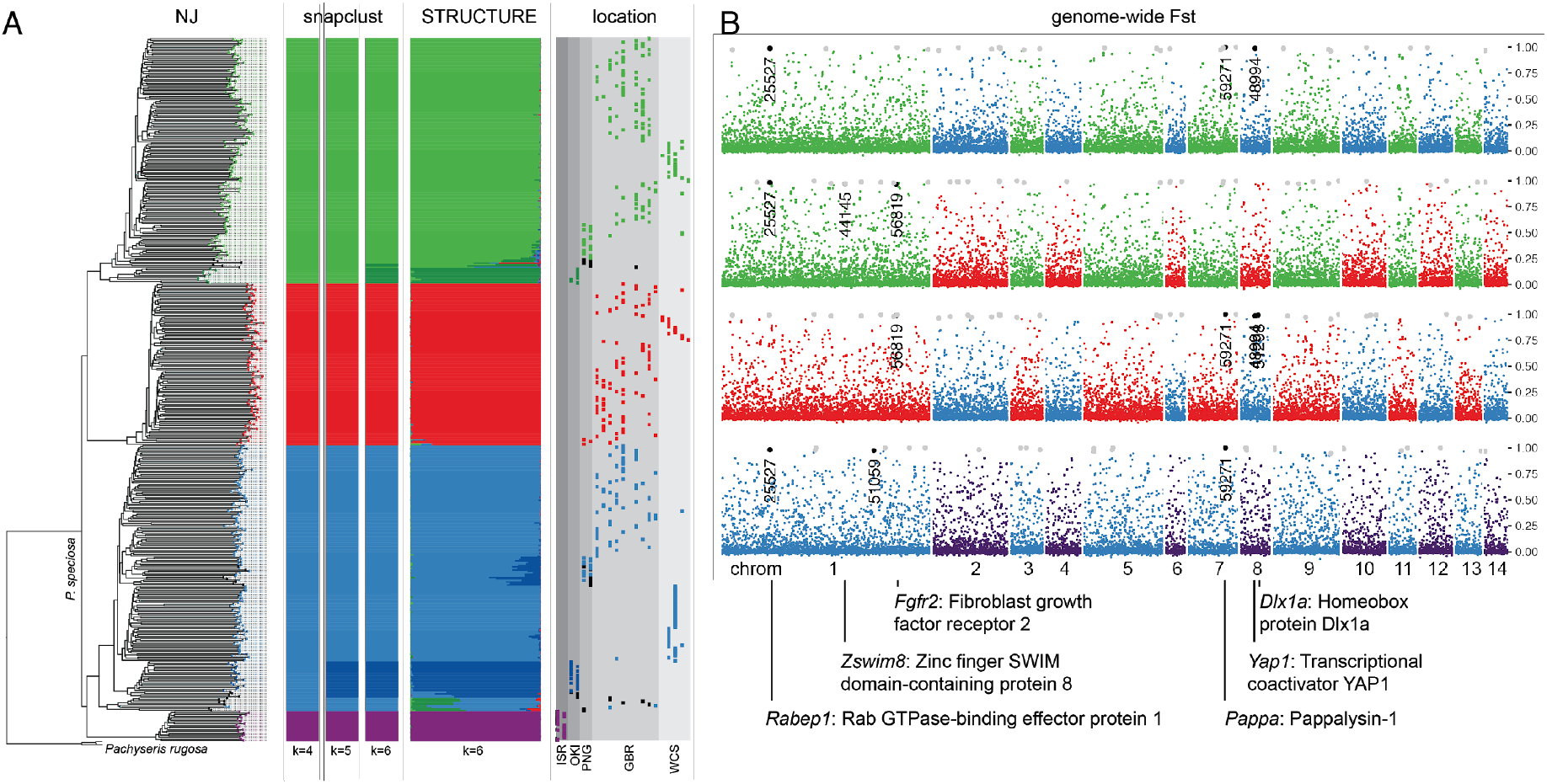
Genetic structuring and genome-wide differentiation of *P. speciosa* lineages. (A) Neighbor-joining tree based on variant nextRAD sites, with matching snapclust (k = 4 to 6) and STRUCTURE (k = 6) assignments for each coral colony. Dots in the “location” column indicate sampling population (geographically arranged within region) with the color indicating STRUCTURE assignment (also indicated in the branch tips of the tree; black indicates potentially admixed samples with a maximum assignment of < 0.95). The bottom clade represents the *Pachyseris rugosa* outgroup (n = 3) and is not included in the adjacent clustering analyses. (B) Manhattan plots showing genome-wide distribution of FST values for different *P. speciosa* lineage comparisons. Individual single-nucleotide polymorphisms (SNPs) are plotted along pseudochromosomes (mapped to *Acropora millepora* for visualization purposes; showing only those that mapped), with the alternating colors indicating the two lineages compared in each plot (“green”, “red”, “blue” and “purple”). Gray and black dots indicate SNPs that are alternatively fixed, with black SNPs indicating those that are missense variants (labelled with their gene ID). Missense variants relating to the inter-comparison of the three Australasian lineages are additionally labelled with the corresponding UniProt gene and protein names (note that two variants, identified as located in *Hecw2* and an unidentified gene, are not mentioned here as they did not map to *A. millepora* chromosomes).

Compared to the divergent, congeneric species *Pachyseris rugosa* [characterized by a more irregular and bifacial morphology (2), outgroup at the bottom of the tree; Figure 2a], the three *P. speciosa* lineages in Australasia represent a closely-related species complex. Differentiation between the “red”, “green” and “blue” lineages is observed across the genome and global mean FST values are much higher (average = 0.1256; range: 0.1032-0.1613) than between geographic regions within these lineages (average = 0.0207; range: 0.0130-0.0339) (Figure 2, S4). The two lineages in Okinawa are more related to the “green” and “blue” lineages, respectively, from Australia than to each other (global mean Fst = 0.1324). Similarly, the “purple” lineage exclusive to Israel (Red Sea) is on average the most divergent of all the lineages, with the closest relatives representing the “blue” lineage from Australia. The apparent restricted distribution of the “red” lineage in Australasia indicates that it may have originated in that region. Indications of admixture (i.e. samples with a maximum assignment < 0.95) between the three lineages were minimal with the exception of one F1 hybrid on the Great Barrier Reef (Figure 2, S5). This coral had a roughly equal assignment to the “green” and “blue” cluster, and a heterozygous genotype for 35 out of 39 genotyped SNPs that were alternatively fixed between corals belonging to these two clusters. Admixture signatures of most other putatively admixed samples were indicative of analytical artefacts (e.g. due to admixture with an unsampled population/lineage; Figure S5). Overall, the limited admixture and presence of only a single F1 hybrid (with its reproductive viability being unknown), indicates that the three lineages have evolved reproductive isolation. This, combined with their widespread sympatric distribution, confirms that these lineages represent distinct species (with their formal description underway; Muir and Bongaerts in preparation), adding to the ever-growing number of cryptic species within the Scleractinia (21, 26).

### Morphologically cryptic lineages despite ancient divergence

An assessment of *in situ* photographs of colonies from the Great Barrier Reef and Western Coral Sea unveiled extensive morphological variability within lineages, but with no apparent distinguishing features between lineages (n = 157; Figure S6). Examination of qualitative traits (including known diagnostic traits for the genus) using skeletal specimens, again found substantial variation but this was not partitioned across the three different lineages (n = 36; Table S9). Although some of the skeletal features overlapped with those described for other *Pachyseris* species (2, 42), the lineages did not align with one of the other five taxonomic species described in this genus [of which only two are reported from Australasian waters; (2)]. An initial assessment of micro-skeletal features using scanning electron microscopy also did not reveal differentiating characteristics between the three lineages (n = 15; Figure S7).

In contrast to the lineages being morphologically indistinguishable, they are divergent genomically with 146 alternatively-fixed SNPs among the three lineages in Australia (0.5 % of 29,287 SNPs; Figure 2). This included 8 missense variants (i.e. a nonsynonymous substitution producing a different amino acid) located in genes of which 7 had homology to UniProtDB genes (IDs: *Rabep1*, *Yap1*, *Pappa*, *HecW2*, *Zswim8*, *Fgfr2* and *Dlx1a*). The Fibroblast growth factor receptor 2 gene (*Fgfr2*) is thought to be involved in environmental sensing and larval metamorphosis (43–46). The *Dlx1a* gene is part of the homeobox-containing superfamily generally associated with morphogenesis (47). The *Pappa* gene encodes the pregnancy-associated plasma protein-A, which is upregulated during spawning in *Orbicella franksi* and *Orbicella annularis* (48).

Whole-genome resequencing of representative “green”, “blue” and “red” colonies from the Great Barrier Reef (n = 20 at 4-29X coverage; Table S10) further confirmed the strong genetic differentiation with minimal admixture (Figure 1f, S8). To assess whether the three lineages have diverged recently and/or experienced differences in demographic history, we used the Pairwise Sequentially Markovian Coalescent [PSMC; (49)] approach for a subset of colonies that were sequenced at greater depth (14-27x coverage; n = 10). Modelled demographic histories showed much greater variation between than within lineages, suggesting an ancient divergence as far back as 10 mya (Figure 3), and that the cryptic nature is due to morphological stasis rather than recent divergence. All lineages peaked in effective population size around 1-3 mya which is consistent with PSMC analyses in recent studies on other robust (50) and complex (51) corals. We also constructed circularized mitochondrial genomes from the whole-genome resequencing data (~19 kbp in length), which in contrast only showed a limited assortment of mitochondrial haplotypes into lineage-specific groups (Figure S9). Given the limited nuclear admixture, this is likely to reflect incomplete lineage sorting and/or ancient introgression, combined with very low mutation rates typical of anthozoans (52, 53). It also reiterates how mitochondrial regions in corals for species-level phylogenetics should be used cautiously (39).

**Figure 3.**
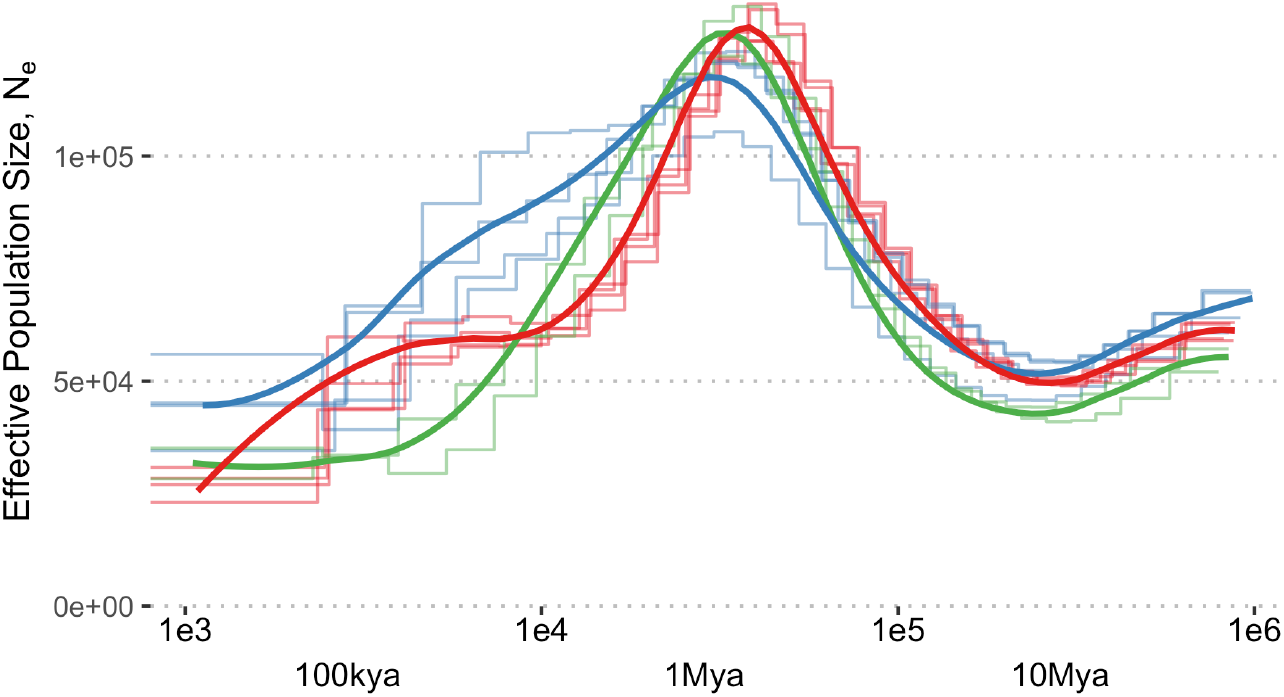
Demographic history as inferred through whole-genome resequencing. Changes in inferred, effective population size as estimated using MSMC for 10 deeply sequenced coral colonies representing the three *Pachyseris* lineages occurring sympatrically in Australia. The timescale is shown in units of numbers of generations in the past. Additional x-axis labels show time in the past assuming a generation time of 35 years. A mutation rate of 4.83e^-8^ was assumed for all calculations (50).

### Ecological and phenotypic differentiation of the lineages

Given the lack of diagnostic morphological features discriminating the three *Pachyseris* lineages, we designed a cleaved amplified polymorphic sequence (CAPS) assay based on the nextRAD data to allow for high-throughput or field-based genotyping (Figure S10; Table S11). We used the rapid assay to expand the number of identified genotypes in Australia (n = 1,442) to include a broad range of reef habitats and locations, and assessed whether there are other ecological and/or phenotypic traits that distinguish the three lineages. The genotyping confirmed the ubiquitous sympatric distribution, with 39 out of 45 sampled Australasian sites containing representatives of all three lineages (Figure 4A). However, there was a significant overall effect of habitat and region on the relative abundances of the three lineages, with pairwise tests indicating significant differences between shallow (back-reef and 10 m) and mesophotic (40 m and 60 m) habitats (confined to GBR and WCS; Table S12; Figure 1G). When assessing the proportions of individual lineages over depth, the “red” lineage was significantly more abundant on shallow (back-reef and 10 m) as compared to intermediate (20 m) and mesophotic (40 and 60 m) habitats, with the opposite pattern observed for the “green” and “blue” lineages (Figure 4b; Table S13). Although the “red” lineage was not found in Okinawa, the divergent “green” and “blue” lineages (as identified using nextRAD sequencing) appeared to be associated with respectively shallow and mesophotic depths (Figure 1G, 4A), indicating there may also be distinct ecological distributions in Okinawa.

**Figure 4.**
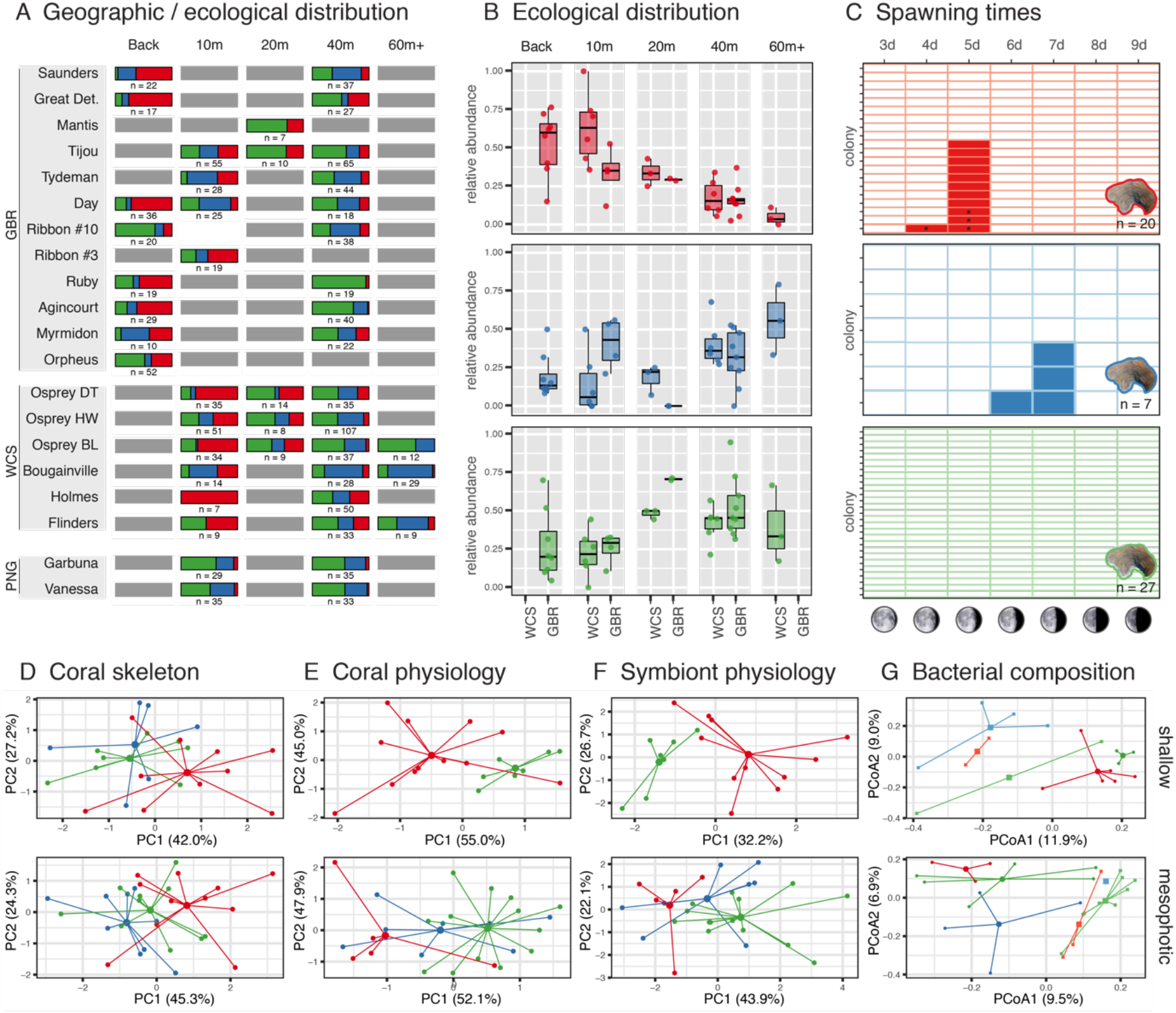
Ecological, phenotypic and reproductive characterization. (A) Proportional composition of *Pachyseris* lineages across geographic locations and depth/habitat. Assignments are based on the CAPS genotyping assay merged with the nextRAD data. Locations within regions are ordered from north to south (top to bottom). (B) Boxplots summarizing relative abundances of *Pachyseris* lineages for each habitat and region (Great Barrier Reef and Western Coral Sea). Dots represent the proportions for each individual population (n = 41; corresponding to those in panel A). (C) Binary heatmap indicating the spawning status of 54 *Pachyseris* colonies (rows) over time (columns) and grouped by lineage. Colonies were collected from shallow depths at Orpheus Island (Great Barrier Reef) and monitored *ex situ* between 3 and 9 days after the full moon in November 2017. Opaque cells indicate the release of eggs (asterisk) or sperm (no symbol). Empty cells indicate that the colony was isolated and monitored, but that no gametes were observed (e.g. no gamete release was observed for the “green” lineage). (D) Principal component analysis for four host skeletal traits. (E) Principal component analysis for two coral host physiological traits: protein and lipid content; PCA). (F) Principal component analysis for six Symbiodiniaceae photophysiological traits. (G) Principal coordinate analysis for bacterial community structure (based on 16S amplicon sequencing). Coral colonies are compared within their own depth groups: shallow (top plots; 10-20 m) and mesophotic (bottom plots; 40-60 m), with most colonies originating from either 10 or 40 m. Skeletal and physiological traits were measured for samples the WCS, whereas the bacterial included samples from both the GBR (square shapes) and WCS (circle shapes) regions.

To examine potential phenotypic differentiation, we quantitatively assessed skeletal and physiological differences between lineages in Australia. Principal component analyses based on coral host skeletal traits (four traits; n = 54), physiological traits (protein and lipid content; n = 69) and Symbiodiniaceae physiological traits (cell density and five pigment traits; n = 70) revealed separation between lineages despite high levels of variance (Figure 4D-F). The “green” and “red” lineages differed significantly across all three trait groups, whereas the “blue” and “red” lineages differed in skeletal traits and “blue” and “green” in symbiont physiological traits (Table S14). In terms of skeletal traits, the mean density of septa (i.e. vertical blades inside the corallite) was significantly lower in the “red” lineage, but not morphologically diagnostic given the overlapping ranges (Figure S11; Table S15). Observed ranges of physiological traits were similar to those reported in other studies that include *P. speciosa* (34, 35); however separating the three lineages revealed differences in protein density, Symbiodiniaceae density, and photosynthetic pigment concentrations (Figure S12; Table S15). Although *P. speciosa* (*sensu lato*) prefers shaded or deeper environments, it has been considered an efficient autotroph across its entire depth range (36). However, we observed a tendency for different lineages to vary in how they trade symbiont densities for chlorophyll concentration per symbiont cell, where lineages with thicker tissues (greater protein densities) hosted greater symbiont densities (Figure S12; Table S15). Although sample sizes were small relative to the observed variance, the results demonstrate the existence of phenotypic differences between the morphologically cryptic lineages, pointing towards distinct physiological strategies.

To explore whether the phenotypic differences may be attributed to differences in coral-associated bacteria or algal symbionts (Symbiodiniaceae), these microbial communities were compared among the three lineages. Associated bacterial communities were assessed by genotyping the host lineages from samples of a previously published 16S rRNA gene metabarcoding dataset [n = 43; (35)]. No significant differences were found in the bacterial community structure, richness, and diversity of the three lineages (Figure S13), but regional patterns were present (WCS and GBR, Figure 4G, Table S16) confirming earlier findings (37). Three bacterial operational taxonomic units (OTUs) from the genera *Corynebacterium* and *Gluconacetobacter* were consistently found in the three lineages, across regions. Despite the lack of differences in the overall bacterial communities, there were two unique OTUs that were only found in the “blue” and “red” lineage, respectively (Figure S13).

To investigate potential differences in lineage-associated Symbiodiniaceae communities, we screened the nextRAD data for contaminating chloroplast or mitochondrial Symbiodiniaceae loci. We found three organellar loci that were genotyped for several hundred samples, but these were largely invariant within Australian samples. This is in line with a previous study (from West Australia) that observed a single Symbiodiniaceae type in *P. speciosa* over depth (34), although some depth partitioning was observed in the Red Sea (35). More detailed mitochondrial characterization was conducted by aligning whole-genome resequencing data from the three lineages to the *Cladocopium goreaui* genome [C1; (52)]; this revealed a highly reticulated haplotype network based on an 8 kb mitochondrial region with 99.7% similarity across samples (n = 16). Out of the eleven haplotypes, two were shared between the “green” and “blue” lineages, whereas the remaining haplotypes were found only once or multiple times but in a single lineage (Figure S14). Overall, microbial associations appeared to be either primarily driven by environment (in the case of the bacterial communities) or were highly consistent (Symbiodiniaceae), indicating that observed phenotypic differences between lineages were unlikely driven by distinct microbial associations.

### Temporal reproductive isolation

Given the limited admixture between the sympatrically-occurring *P. speciosa* lineages, we monitored the reproductive behavior of shallow colonies from Orpheus Island on the Great Barrier Reef. Previous reports from that location indicated spawning of *Pachyseris speciosa* between 5 and 6 days after the full moon (55). Colonies were collected from the field just after the full moon in November 2017, genotyped using the CAPS assay (which identified 20 “red”, 7 “blue”, and 27 “green” colonies), and monitored *ex situ* for spawning from 3 to 9 days after the full moon. On day 5, half of the colonies of the “red” lineage released gametes (with one colony releasing also on day 4), with no colonies from the other two lineages releasing gametes (Figure 4C). On day 7, nearly half of the colonies from the “blue” lineage released gametes (with one also releasing on day 6; Figure 4c), with again no colonies from the other two lineages releasing gametes. No gamete release was observed for the “green” lineage within the monitoring period, although the “green” colony from which the draft genome was constructed spawned 8 days after the full moon in December 2014 (see Methods). Overall, the average spawning time of male colonies was 19 min after sunset (n = 13), versus an average of 43 min for female colonies (n = 4; Table S18), in line with general observations of males spawning earlier in broadcast spawning coral (56). Attempts to preserve unfertilized eggs in filtered seawater (at ambient temperature) for experimental inter-lineages crosses were not successful (complete degradation within 24 hours).

The temporal segregation in the timing of gamete release observed during the November 2017 spawning provides a potential mechanism to explain the minimal admixture in these sympatrically-occurring lineages. Temporal reproductive isolation is a common strategy for minimizing interspecific gamete encounters in scleractinian corals, with the timing difference ranging from hours (57, 58) to months (59, 60). The two-day difference in gamete release between “red” and “blue” *P. speciosa* points towards differences in processes entrained by the lunar cycles, rather than environmental seasonal or diurnal cycles (61–63). The identification of fixed high-impact gene variants related to environmental sensing, development, and gametogenesis provide initial leads to the genomic basis of this potential prezygotic reproductive barrier.

Given the predicted split-spawning (i.e. mass spawning occurring across at least two consecutive months) in 2017, we maintained a subset of the colonies (n = 36) for additional spawning monitoring after the full moon of December in that year. Observations undertaken for those colonies saw gamete release for only one colony of the “blue” lineage and one of the “green” lineages (both on day 4 after the full moon). In total, two-thirds of the colonies from the “red” lineage released gametes, with release observed in the period between day 3 and day 8, with most of the release occurring on day 4 (8 colonies versus 1-3 colonies on the other days) (Table S17). While these observations confirmed the occurrence of a split spawning, they may not reflect natural release patterns, given the extended time these colonies were removed from natural cues (e.g. exposure to lunar cycle, lack of exposure to tides). In addition, their prolonged close proximity to one another in the single tank (“raceway”) in which they were kept, may have affected the timing of release [e.g. due to chemical signaling; (62)]. The observations confirm the potential for experimental inter-lineage crossings under artificial conditions, to assess whether temporal reproductive isolation is accompanied by gametic incompatibility.

### Geographic and habitat structuring within lineages

Within each of the three Australasian lineages, principal component analyses identified considerable substructuring, which was driven to a large extent by sampling region (Great Barrier Reef, Western Coral Sea and Papua New Guinea; Figure 5A,C). However, additional substructure was identified (2-3 apparent clusters) in the GBR samples that could not be linked to subregions, locations or habitat (Figure 5C). Discriminant analysis of principal components (DAPC) further confirmed the genetic differentiation among the three sampling regions (Figure 5B), indicating restricted gene flow between these regions with a some exceptions indicating recent admixture. Although genetic differentiation between the GBR and WCS has been observed previously in other coral species (64–65), here we identify further differentiation between individual WCS atolls confirming their rather isolated nature as they are surrounded by deep oceanic water. In contrast, only very subtle substructuring was identified between the “Northern” and “Far Northern” regions of the GBR (Figure 5C). The inability to detect distinct location- or region-associated clusters on the GBR could indicate the potential for gene flow over large distances, a pattern observed for other coral species and facilitated by the high density of reefs along the outer shelf (64–68). Nonetheless, given the presence of substructuring within each of the locations [a pattern frequently observed in scleractinian corals; (22)], such “panmixia” should be interpreted with caution.

**Figure 5.**
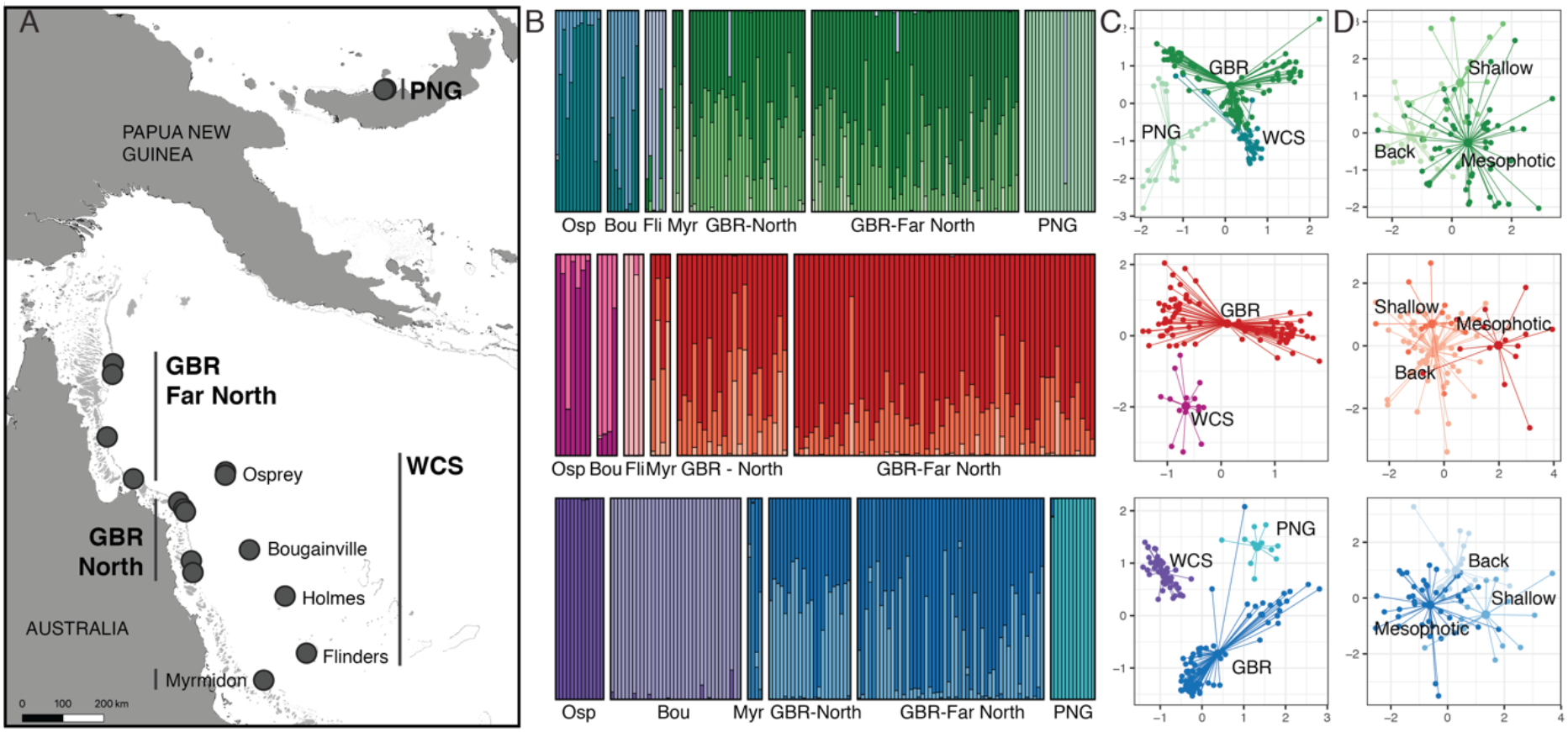
Genetic structuring within lineages by region and habitat. (A) Map showing the Australasian sampling locations/regions included in the DAPC plots. (B) DAPC assignment plots for each of the three lineages (from top to bottom: “green”, “red” and “blue”) using location as prior (but ignoring habitat and grouping “North” and “Far North” locations on the Great Barrier Reef). (C) Principal component analyses for each of the three lineages (from top to bottom: “green”, “red” and “blue”), showing structuring by region as well as unexplained substructuring within the Great Barrier Reef populations. (D) DAPC scatter plot for the Great Barrier Reef using habitat as prior. PCA and DAPC scatter plots show individual colonies connected to respectively region and habitat centroids, and analyses are based on nextRAD data after the removal of outliers.

Where other studies found a clear partitioning of cryptic lineages across habitats (28, 29), we observed substantial overlap between the depth distributions of the three *Pachyseris* lineages (e.g. the “red” lineage is present at shallow and mesophotic depths, but at lower relative abundances in the latter). Surveys during the 2016 mass bleaching found that most *Pachyseris* colonies down to 25 m were bleached, while fewer than half of the colonies at 40 m depth were bleached (69). Opportunistic genotyping of 14 healthy and 3 bleached colonies collected from 40 m during this event found healthy representatives of all three lineages (8 “green”, 5 “red”, and 1 “blue” healthy colony; versus 2 “green” and 1 “blue” bleached colony). The broad depth range of these lineages combined with the strong depth-attenuation of bleaching in *Pachyseris* (69), raises the question whether deep populations may benefit those at shallow depth by acting as a refuge and source of reproduction (70). Unfortunately, genetic differentiation between habitats (within each lineage) could not be adequately assessed at individual locations given the small sample sizes after splitting into cryptic lineages. When merging individual locations on the GBR, DAPC showed some discrimination between habitat clusters (Figure 5D, S15). This separation was driven by many low-contributing and several high-contributing alleles indicating there may be potential limitations to such vertical connectivity (Figure S15).

## Conclusions

Tens of thousands of multicellular species have been described in association with tropical coral reefs, yet this is estimated to represent <10% of their actual species richness (1). Reef-building corals (Scleractinia) have been considered to contribute relatively little to this estimated discrepancy, with the taxonomy considered to be relatively complete compared to other less-studied coral reef taxa (1). Molecular approaches have indeed corroborated many of the current taxonomic species (71), but have also unveiled extensive undescribed diversity within the traditional morphological boundaries (21, 26). This pattern is likely to be accelerated given the transition to genome-wide sequencing approaches, solving the resolution issues associated with traditional sequence markers (30, 72, 73). The inability to discriminate this diversity as readily identifiable taxonomic units in the field (or even through close skeletal examination) has raised concerns about the practicality and necessity of incorporating it into our systematic framework, particularly as the lack of morphological differentiation can lead to the assumption of negligible physiological differentiation. While we found no diagnostic morphological characters distinguishing the three *Pachyseris* lineages in this study, the genome-wide differentiation was accompanied by differences in ecological, physiological, and reproductive traits, demonstrating how morphological stasis can mask substantial ecological divergence. Failure to recognize such cryptic diversity will result in erroneous interpretations of species distributions, extinction risk, and spatial genetic differentiation (21, 26), all with critical ramifications for conservation management. In light of the rapid degradation of the world’s coral reefs, it is critical to acknowledge and start capturing this hidden diversity to improve our understanding and ability to protect these fragile ecosystems.

## Methods overview

We assembled and annotated a *de novo* reference genome using PacBio sequencing (~100x coverage) of a sperm sample from a *Pachyseris speciosa* colony from Orpheus Island on the Great Barrier Reef. This was used as a reference for reduced-representation sequencing (nextRAD) of *P. speciosa* colonies (n = 501) from shallow and mesophotic habitats in five different regions (Great Barrier Reef, Western Coral Sea, Papua New Guinea, Okinawa and Israel), sequenced across 4 Illumina NextSeq 500 lanes. Whole-genome re-sequencing of representative *P. speciosa* samples (n = 20) was then undertaken for historic demographic modelling, using the Illumina Nextera protocol at ~5X or ~20X coverage across 8 Illumina HiSeq 2500 lanes. Given the lack of diagnostic morphological characteristics distinguishing the lineages, we designed a cleaved amplified polymorphic sequence (CAPS) assay to increase the number of genotyped samples (n = 1,442), and assess ecological, phenotypic and reproductive differences. Morphological differences between lineages were assessed through *in situ* photographs (n = 157), examination of qualitative traits in bleach-dried skeletons (n = 36), and micro-skeletal features using scanning electron microscopy (n = 15). Quantitative measurements were undertaken for five skeletal characters (n = 54). Physiological characterization was undertaken through protein and lipid quantification, *Symbiodiniaceae* cell counts, and photopigment quantification using high-performance liquid chromatography (HPLC) (n = 73). Spawning behavior was assessed through reproductive monitoring of colony fragments (n = 54) during November/December 2017 at the Orpheus Island Research Station. Microbiome characterization was undertaken by genotyping host lineages (n = 43) of a previous study based on 16S rRNA amplicon sequencing (37). See *SI Appendix, Supplementary Methods* for a detailed explanation of all the methods.

## Data Availability

Raw sequence data for the genome and transcriptome assembly are available through ENA accession number <u>PRJEB23386</u>, the reduced-representation data through <u>XXXX</u>, and the whole-genome resequencing through <u>XXXX</u>. Genome assembly, variant call datasets, electronic notebooks, and scripts are accessible through https://github.com/pimbongaerts/pachyseris.

## Acknowledgements

We thank David Whillas, Jaap Barendrecht, David Harris, Sara Naylor, Annamieke van den Heuvel for support in the field or in the lab, as well as Underwater Earth, The Ocean Agency, and crews from Reef Connections, Mike Ball Dive Expeditions, SY Ethereal and the Waitt Foundation. The authors acknowledge the Reef Future Genomics (ReFuGe) 2020 Consortium, of which this study was part, as organized by the Great Barrier Reef Foundation.

## Author contributions

PB, IRC, HY, DW, SdH, MAR, MJHvO, OHG conceived and designed the research. PB, DW, SdH, CAB, SD, NE, GE, MG, KBH, DCH, MM, PM, GT contributed to lab work. PB, NE, GE, KBH, SH, AM, FS, PSW, GT contributed to specimen collections. PB, IRC, YH, DW, SdH, AHA, SF, YL contributed to analyses. PB wrote the paper, and all authors contributed manuscript edits.

