## Supplemental Information for "Morphological stasis masks ecologically divergent coral species on tropical reefs"

### "Cryptic diversity masks ecologically distinct coral species on tropical reefs" - SUPPLEMENTARY

1. Methods 1
   1.1 Specimen collections 1
   1.2 Genome assembly and annotation 2
   1.3 Reduced-representation sequencing 3
   1.4 CAPS marker development and genotyping 5
   1.5 Whole-genome re-sequencing and demographic analyses 6
   1.6 Characterization of associated Symbiodiniaceae and microbiomes 7
   1.7 Morphological and physiological characterization 8
   1.8 Reproductive characterization 9
2. Supplementary tables 10
3. Supplementary figures 21
4. References 34

#### 1. Methods

##### 1.1 Specimen collections

For the reference genome, sperm from a single colony of *Pachyseris speciosa* was collected at Orpheus Island Research Station (under GBRMPA permit G14/36802.1). The colony was collected on 8 December 2014 from the Island's fringing reef (S18.608°, E146.489°) from 6 m depth, and kept in a flow-through aquarium with 0.5 micron filtered seawater. Every afternoon, one hour before sunset, the colony was placed in a container with as little filtered seawater as possible to cover the whole colony. Broadcast spawning of sperm was first observed on 14 December, 2014 at 18:35 (exactly at sunset). The colony's holding water was then centrifuged in 50 mL falcon tubes to concentrate the sperm into a pellet, using an Eppendorf 5702 R centrifuge at 3,000 g for 15 min. The resulting sperm pellet was snap-frozen in liquid nitrogen and stored at -80 °C. Additional tissue from the colony was sampled on 15 December, 2014, and stored at -80 °C for transcriptome sequencing. For population-level assessments, small fragments from a total of ~2,500 *P. speciosa* colonies were collected (details in Table S1) from the Great Barrier Reef and Coral Sea atolls in Australia, Kimbe Bay in Papua New Guinea, Okinawa in Japan, and Eilat in Israel. Primary collections were performed across three distinct reef habitats: the back-reef (10 m ± 3), shallow slope (10 m ± 3) and deep slope (40 m ± 3), with additional populations collected at intermediate (20 m ± 2) and lower mesophotic depths (60-85 m). Samples were collected using SCUBA or a remotely operated vehicle (ROV; Seabotix vLBV300) between 2012-2017 (with the vast majority collected between September-December 2012). Small fragments of the collected specimens were stored in salt-saturated buffer solution (containing 20 % DMSO and 0.5 M EDTA) and/or in molecular-grade ethanol, and for a proportion of specimens a skeletal voucher was bleached, rinsed in freshwater and dried. Species distribution map of *P. speciosa* was downloaded from the IUCN Red List of Threatened Species (1).

##### 1.2 Genome assembly and annotation

High-molecular-weight DNA was extracted from sperm using a method based on Blin and Stafford (2). Initial sequencing was undertaken on an Illumina HiSeq 2500 following the PCR-free library construction protocol developed by the Broad Institute. For the genome assembly, long-read sequencing was then undertaken on the PacBio Sequel platform at the Ramaciotti Centre for Genomics, using across 92 SMRT cells to ~100x coverage. RNA was extracted from tissue collected from the same colony, with directional RNA-seq libraries sequenced on an Illumina HiSeq 2500 at the Australian Genome Research Facility (AGRF). Prior to the assembly, genome size and heterozygosity rate were estimated based on the paired-end short reads. FastQC (3) was first applied to screen the base quality, GC-content, overrepresented k-mers, and adaptors. Genome size estimates were calculated using sga.preqc (4) and GenomeScope (5) based on a k-mer size of 31, and were respectively 886.1 and 749.6 Mb (Table S2).

The final genome assembly was achieved in three stages. Firstly, the genome assembly was conducted using CANU v1.5 (6) with default settings. This resulted in 10,783 contigs, 1,788 Mb assembled sequences and N50 size of 328 Kb. This is in line with the observation that the assembled size is nearly twice as large as the true genome size when the heterozygosity rate is high in a diploid genome (7). To reduce allelic redundancy, HaploMerger2 [version 20161205; (8)] was then employed, which successfully merged 44% of the sequences. Finally, PacBio long reads were mapped to the assembly using BLASR (9) and the mean coverage was calculated for each contig. Contigs whose GC contents were greater than 45% (genome GC% = 39%) and read coverages were lower than 50X (~ half of the expected coverage) were considered as putative contaminants. These contigs were used as queries to perform blastn search (E-value $\leq 1e^{-05}$) against the NCBI non-redundant nucleotide database (10). Contigs that contained more sequences significantly similar to non-metazoan sequences than to metazoan sequences were removed.

*De novo* identification of repeat classes was accomplished with RepeatModeler v1.0.11 (11) with parameter "-engine ncbi". The classifier utilized two *de novo* repeat finding programs, RECON and RepeatScout, and was built upon RepBase v20181026 (12). The resulting repeat library was used as input by RepeatMaster v4.0.8 (10) to generate the repeat annotation. Protein-coding gene annotation was performed as described previously (13). Briefly, de novo and genome-guided transcriptome assembly was performed using Trinity (14), followed by PSyTrans to remove Symbiodiniaceae transcripts (15). Transcripts were then assembled to the genome assembly using PASA (16), from which a set of likely ORFs were generated. Based on their protein coding ability and completeness, these ORFs were carefully assessed to produce a high confidence and non-redundant training gene set. This was used to train AUGUSTUS (17) and SNAP (18), and the resulting parameters were employed by the corresponding program from MAKER. The *ab initio* gene model was predicted using the MAKER2 (19) pipeline. In addition, putative transposable elements in the gene model were excluded based on transposonPSI (20) and hhblits (21) searches to transposon databases.

Both the genome assembly and gene model datasets were tested for the completeness of conserved core genes using Benchmarking Universal Single-Copy Orthologs (BUSCO) software (22). BUSCO program version 3 was run with default parameters. The metazoan gene set (odb9), which contains 978 orthologs, was employed as the reference dataset. To further validate the gene model, the predicted protein sequences were matched against the Swiss-Prot database and PFAM-A protein domain database. Swiss-Prot database (2018-08) was downloaded from UniProt FTP site (23) and blastp was performed (E-value ≤ $1e^{-05}$). The annotated coral proteins were used as queries and the curated database proteins were used as targets. The target coverage was defined as the percentage of the target length in the alignment. To identify well-defined protein domains, HMMER [hmmer3 (24)] was used to perform alignments to PFAM-A (v31.0) hmm profile, and those with a combined E-value and c-E-value lower than $1e^{-05}$ were selected.

##### 1.3 Reduced-representation sequencing

Coral gDNA extraction was performed as in Bongaerts et al. (25), reducing *Symbiodiniaceae* contamination through several centrifugation steps, unless this resulted in insufficient gDNA yield (<150 ng gDNA) in which case the extraction was performed directly on the sampled tissue (~25% of sequenced samples). Quality and yield of gDNA were assessed using gel electrophoresis and a Qubit fluorometer to select a subset of higher-quality samples within each sampled population for downstream sequencing (giving preference to those where endosymbiont contamination was reduced; n = 672). This included 3 replicates of the same sperm sample used for the reference genome, 7 additional technical replicates, and a *Symbiodiniaceae* sample isolated from a *P. speciosa* colony using fluorescence-activated cell sorting [following the same protocol as in Bongaerts et al. (25)]. Library preparation was carried out using the nextRAD method (Nextera-fragmented, reductively-amplified DNA; SNPsaurus, LLC), which uses a selective primer sequence (rather than restriction enzymes) to genotype loci consistently between samples. Genomic DNA was purified using AMPure XP beads, and then fragmented and ligated with adapter sequences using Nextera reagent (Illumina, Inc). Fragmented DNA was then PCR amplified (73 ºC for 26 cycles) with one of the primers matching the adapter and extending into the genomic DNA using a 9 bp selective sequence ("GTGTAGAGG"). Libraries were sequenced across 4 NextSeq 500 (Illumina, Inc) lanes using 150 bp single-end chemistry and following the manufacturer's recommended protocol. Parsing and analyses are detailed in an electronic notebook (26), using generic Python scripts located in a separate repository (27). Most of the statistical analyses and plotting was performed in R (v3.5.3; R Core Team 2019).

TrimGalore (28) was used to trim Nextera adapters and low-quality ends (PHRED < 20), while discarding reads shorter than 30 bp. Reads of each sample were then mapped to our *P. speciosa* genome using BWA-MEM (29) as well as *Cladocopium goreaui* genome [ITS2 type C1; (30)], with the latter to assess overall sample performance and contamination (for the two extraction methods and different gDNA yields; Figure S1). Variant calling of the resulting BAM files was done using "UnifiedGenotyper" from the GATK pipeline (31), and then hard-filtered using "VariantFiltration" for bi-allelic single-nucleotide polymorphisms (SNPs) with a minimum coverage of 10X, a minimum genotype quality of 30, and a minimum allele frequency of 0.01. Initially, the overall dataset was reduced to those SNPs that were genotyped for at least 80% of samples and those samples that were genotyped for at least 50% of SNPs (n = 501). Genotyping accuracy was verified from three replicate sperm samples (99.8% similarity), as well as seven regular replicate pairs (separate gDNA extraction and library preparation; 98.9-99.6% similarity), with only the highest-performing sample of each replicate set retained in the eventual dataset. Duplicate genotypes (clones) were identified using the "vcf_clone_detect" script (27), using a conservative manual threshold (<97%) based on the determined genotyping accuracy (from technical replicates) and the distribution of pairwise genetic similarities across all samples (Hamming-based), retaining the sample with the least missing data from each set of duplicates (total of 24 potential clones removed). Given that the opportunistic sampling could have led to accidental resampling of colonies, no interpretations were made regarding clonality rates.

To assess overall genetic structuring, we visualized the structure of the overall SNP dataset (with 468 remaining samples; 8,536 SNPs) using a Principal Component Analysis (PCA) in adegenet (32), and a neighbor-joining (NJ) tree based on genetic distance [Hamming-based; using the Phylo module in the Biopython library (33)]. Both indicated the presence of 4 major clusters in our data (Figure S2), which was then further explored using snapclust (34), a maximum likelihood approach based on the Expectation-Maximisation (EM) algorithm that offers goodness-of-fit statistics. We evaluated three different statistics (Akaike, Bayesian and Kullback Information Criteria) using the "choose.k" function under increasing numbers of clusters (up to 20) across five replicate, subsampled datasets (thinned to ensure a minimum distance of 2,5 Kbp between SNPs). We then ran snapclust for the indicated optimum window (k=4 to k=6; Figure S2), again using the same five replicate datasets, using the "Ward" algorithm to define initial group assignments, and with 50 iterations of the EM. At k=5 and k=6, two additional geographic populations were separated out from the four most divergent clusters, which was also supported in the NJ tree (Figure 2), and k=6 was therefore chosen for subsequent analyses. Using the "structure_mp" wrapper (27) we then also ran STRUCTURE [v.2.3.4; (35)] for 20 replicate, subsampled datasets (again thinned to ensure a minimum distance of 2,5 Kbp between SNPs) to further assess potential signatures of admixture between the 6 clusters. Runs were conducted using the admixture model with correlated allele frequencies and not considering priors (50,000 repeats after a burn-in of 100,000). Individual runs were then aligned using CLUMPP [v.1.1.2; (36)], and assessed for the presence of "ghost clusters" (i.e. clusters with no fully assigned samples), only retaining those runs that have a maximum ancestry assignment of at least 0.99 across all clusters. Samples were then separated out to clusters using a "lenient" ($\geq 0.8$) and "stringent" ($\geq 0.95$) mean ancestry assignment cut-off (marking samples below those thresholds as "unassigned"). The assignment was compared to clustering in the NJ tree by coloring branch tips according to their "lenient" STRUCTURE assignments (Figure 2). Potential admixed samples were identified by extracting those with a maximum ancestry assignment of $< 0.95$ across clusters. The potential of one specific sample to represent an F1 hybrid was assessed by quantifying heterozygosity for SNPs that were established to be alternatively fixed for the two clusters the sample was assigned to.

To assess the extent and nature of divergence between the three lineages, a reduced SNP dataset was generated with only stringently assigned samples, grouped by cluster and geographic region, and retaining only SNPs that were genotyped for at least half the samples in each group. Pairwise genome-wide $F_{ST}$ (37) values were calculated between all groups (45 pairwise comparisons) to assess overall divergence between lineages vs geographic regions using vcftools (Danecek et al. 2011). Highly divergent SNPs were extracted by calculating pairwise allele frequency differentials (AFDs) between the three Australian lineages (using the Great Barrier Reef and Coral Sea regions as "replicates"), and identifying those SNPs that across both regions were alternatively fixed (AFD of $\geq 0.95$ to allow for some genotyping error). Genetic variants were annotated and functional effects predicted using SnpEff (38), with divergent SNPs manually assessed based on their UniProt IDs. For visualization purposes only, we used RaGOO (39) to assemble our 2,368 genomic scaffolds into pseudomolecules by mapping them to a chromosome-level assembly of *Acropora millepora* (40), under the expectation that broad-scale synteny is expected to be conserved [e.g. as demonstrated for *Acropora digitifera* and *Nematostella;* (41)]. We obtained Gene Ontology (GO) terms for all SNPs using the UniProt portal (23), and then used Gowinda (42) to conduct GO enrichment analyses, using "gene" mode (assuming complete linkage disequilibrium between SNPs within a gene) for the alternatively fixed SNP sets using a 1000 bp window upstream and downstream, and 1,000,000 simulations. Assessments of these SNPs for gene ontology (GO) enrichment did not identify significantly enriched GO terms (although admittedly only a small proportion of the overall genome was assessed through the reduced representation sequencing).

Genetic structuring within each of the three Australasian lineages was assessed by splitting the dataset based on the stringent assignment cut-off (0.95), and removing samples below that threshold. These three datasets were filtered for SNPs genotyped for at least 50% of samples (within each lineage), and with a maximum observed heterozygosity threshold of 0.5 (to filter out potential paralogs). For each lineage, we also created a dataset that was more representative of neutral genomic diversity by removing SNPs that were identified as outliers using pcadapt (43) based on q-values with an expected false discovery rate lower than 10%. We used pcadapt as it is generally robust under hierarchical population genetic structure and does not require *a priori* population information, which is important given our nested sampling (region, location and habitat) and low population sizes at the deepest level (habitat) associated with the splitting of the dataset (into three cryptic lineages). For both "neutral" and "overall" datasets, we then assessed overall genetic structuring using principal component analysis [PCA in adegenet; (32)], and evaluated structuring across habitats, locations, and regions using discriminant analysis of principal components [DAPC; (44)].

##### 1.4 CAPS marker development and genotyping

In order to develop a rapid and cost-effective diagnostic assay for three cryptic Australasian lineages, we screened the nextRAD sequence loci for lineage-diagnostic mutations in the sequence motifs of commonly available restriction enzymes. As this was undertaken prior to the establishment of the reference genome, clustering and variant calling of the nextRAD data was first analyzed *de novo* using PyRAD v3.0.66 (45) using a clustering threshold of 88%, a minimum sequence coverage of 6, and a maximum of 4 sites with a PHRED quality below 20. Sites that could be targeted using cleaved amplified polymorphic sequence (CAPS) markers were then identified using a custom script ["pyrad_find_caps_markers.py"; (26)]. An initial host genome assembly and realigned whole-genome resequencing data was then used to manually align potential target loci, so that primers could be designed to flank the loci [using Primer3; (46)], and the presence of additional adjacent restriction sites could be evaluated. After initial amplification, digestion and reproducibility screening for 20 different primer pairs, three markers were selected (named "Pspe-Green-CfoI/HhaI", "Pspe-Blue-HaeIII", and "Pspe-Red-Taqα1"; Table S11) and further tested on samples with a known lineage assignment (based on nextRAD) data to confirm genotyping reliability (3 out of 120 samples showed a different assignment compared to that based on the RAD-seq data).

For the amplications, we used 0.5–1.0 µl of DNA, 1 µl 10x PCR buffer (Invitrogen), 0.3 µl 50 mM MgCl2, 0.2 µl 10 mM dNTPs, 0.5 µl for both the forward and reverse primer (10 µM), 0.07 µl of Platinum *Taq* DNA Polymerase (Invitrogen) and $dH_20$ water to a total volume of 10 µl per reaction. The cycling protocol was: 1×94°C (4 min); 31×[1 min at 94°C, 1 min at 61°C, 1 min at 72°C]; 1×72°C (6 min). The restriction digest mix contained 1 µL enzyme buffer (10X), 0.1 or 0.05 µL restriction enzyme depending on the concentration (10.000 u/mL or 20.000 U/mL), 0.5 µL PCR product, and $dH_20$ water to a total volume of 10 µl per digest. The restriction digest was run for 1 hour at 37°C (for CfoI, HhaI, or HaeIII) or 65°C (for TaqαI), and visualized by running 4 µl of digest product on a 3% agarose gel with GelRed® stain (120V for 40 min) immediately afterwards. Field assays were performed using the miniPCR™ thermal cycler and blueGel™ visualization system (Amplyus, Cambridge, MA, USA), using agarose gels with TBE buffer.

Overall, 1,119 samples were genotyped using the assay (1-3 CAPS markers per sample), including those for the physiological, microbial, and reproductive assessments in this study, and together with the RAD-seq data this led to a total of 1,442 genotyped samples across regions. To assess ecological distributions in the Australasian region (n = 1,312; only considering populations with at least 7 samples), we used PERMANOVA as implemented in the adonis function of the R vegan package (47) based on Bray-Curtis dissimilarities. We tested for differences in relative proportions of the three lineages between habitats and regions (considering only the 10 m and 40 m habitats as those were sampled across all three Australasian regions) using location as strata, followed by pairwise testing of all habitats within each region (with *p*-values adjusted using Bonferroni correction for multiple comparisons). To assess differences in the relative abundances across habitats (for each lineage separately), we used one-way ANOVA (Type II) testing for the GBR and WCS regions, followed by pairwise testing (Tukey's test and *p*-values corrected with single-step method) between habitats. Assumptions of normality and homoscedastic were fulfilled (a square root transformation was applied to the relative abundances of the "green" lineage).

##### 1.5 Whole-genome re-sequencing and demographic analyses

Representative samples (n = 20) of the three distinct lineages (as identified by the nextRAD sequencing) were selected for whole-genome re-sequencing (see Table S12). These samples originated from back-reef habitats (10 m) in three regions on the Great Barrier Reef (Central GBR: Myrmidon Reef, Northern GBR: Ribbon Reef 10, and Far Northern GBR: Great Detached Reef). Genomic DNA were purified using AMPure XP beads (1.8:1 beads/DNA ratio), with separated barcoded libraries prepared for each individual using Illumina's Nextera kit following the manufacturer's recommended protocol. Pooled libraries were sequenced on four Illumina Hiseq 2500 (Illumina, Inc) lanes using 100 bp paired-end chemistry resulting in an average coverage of ~5X. Four additional lanes of sequencing were conducted for ten representative samples (representing the three different lineages from both Myrmidon Reef and Great Detached Reef) to obtain higher coverage (~20X) for demographic analyses. All commands used to analyze whole genome resequencing data and perform demographic analyses are detailed in an electronic notebook (48).

Raw reads were first pre-processed and mapped against the *P. speciosa* genome based on GATK best practices (31). Adapters were marked using Picard (v2.2.1), mapping was performed using bwa mem [v0.7.17; (29)] marking shorter split hits as secondary but with all other parameters at their defaults, and PCR duplicates were marked using Picard. Since low coverage samples had insufficient depth to reliably call genotypes, we used a genotype likelihood approach to investigate genetic structuring and assess admixture at the whole genome level. To support this approach ANGSD [v0.913; (29)] was used to call SNPs and calculate genotype likelihoods. Variant sites (SNPs) were retained if they had a p-value less than $1e^{-06}$ (GATK probability model), and a minor allele frequency greater than 5%. Genotype likelihoods called using ANGSD were used to explore genetic structuring and admixture using PCAngsd [v0.973 (51)] which confirmed the existence of three lineages of this study (Figure S8) and also provided estimates of admixture between these lineages (Figure 1F).

Demographic histories for each of the ten deeply sequenced colonies were inferred based on the distribution of heterozygous sites using the PSMC' method (52) implemented in msmc2 (v1.1.0). In order to avoid known inaccuracies due to fragmentation (53) or miscalled genotypes the analysis was restricted to scaffolds larger than 1Mb (a total of 390Mb) and genomic regions with repeats, excessively low or high read coverage were excluded. Demographic histories for each colony were inferred by performing 100 bootstrap replicates and taking the average. Bootstrap data was generated by randomly sampling the genome in 0.5Mb chunks and arranging these into 30 scaffolds of length 20Mb per replicate. Since this approach allows inference of historical effective population sizes for each sequenced coral colony (52), it was possible to assess the consistency of these estimates within lineages. In order to translate msmc results into real timescales and effective population sizes the spontaneous mutation rate, $\mu$ and generation time, g are required. In the absence of data required to independently calculate these parameters we used estimates ($\mu$=4.83e-

8; g=35y) recently published for *Orbicella* (54) as these are the closest available in a phylogenetic sense. It should be noted that there is considerable uncertainty in these estimates which affects the timescale and effective population sizes, however the shape of curves shown in this figure are unaffected by changes in these parameters.

A reference mitochondrial genome for *P. speciosa* was assembled using deeply sequenced data from a single colony from the "red" lineage, sampled at Myrmidon back reef (Central GBR). Mitochondrial reads were extracted from whole genome data, assembled and scaffolded into a single contig of length 19,507 bp using MITObim [version 1.9; (55)] with the complete mitochondrial genome of *Acropora digitifera* (Genbank accession NC_022830) as a bait. Manual inspection revealed overlapping sequence at both ends indicative of a circular sequence. After trimming redundant bases a circular genome of length 19,007 bp with a single gap of length 30 bp was produced. Annotation of this genome was performed with MITOS (56) and the sequence was submitted to Genbank (accession XXXX).

Consensus mitochondrial sequences for each of the whole-genome sequenced colonies were generated by mapping raw reads to the reference mitochondrial genome. Mapping was performed with bwa mem and resulted in a high coverage (>3000x) bam file for each colony. Consensus sequences (n = 20) were then called using samtools (version 1.6) and bcftools (version 1.9) (57). These were imported into Geneious [version 11.0.2; (58)] as an alignment and trimmed to remove a 468 bp region where manual inspection of read coverage suggested a potential misassembly or unresolved repeat. The trimmed alignment was exported to nexus format and visualized with PopArt [version 1.7; (59)] as a TCS (60) network (Figure S9).

##### 1.6 Characterization of associated Symbiodiniaceae and microbiomes

Basic characterization of associated Symbiodiniaceae was done by extracting "contaminant" nextRAD loci matching the plastid genome of *Cladocopium goreaui* [C1; (30)]. For this, all loci from the *de novo* analyzed dataset (constructed with PyRAD for the CAPS development) were mapped against the *C. goreaui* plastid and mitochondrial genomes using bwa mem (29), retaining only loci that were genotyped for at least 100 samples. The loci were visually assessed, and the two most informative nextRAD loci were extracted (both matched a single plastid genome scaffold) and evaluated. We also mapped the Nextera whole-genome re-sequencing to the *Cladocopium goreaui* mitochondrial genome using bwa mem (29), then called the consensus sequence for a 7 kb high-coverage region of 16 *P. speciosa* samples using samtools [version 1.6; (57)], and assessed haplotypes using PopArt [version 1.7; 58)] (Figure S14).

Microbiome characterization was carried out in Hernandez-Agreda et al. (61), and we now determined the host genotypes for 43 of those samples and reanalyzed the data for lineage-associated patterns. Detailed methods are described in Hernandez-Agreda et al. (60). In short, DNA was extracted using the modified protocol (62) of MoBio PowerPlant Pro DNA isolation kit, with the bacterial 16S rRNA gene PCR amplified using the 27F/519R (v1-v3 region) primers. Library preparation was carried out using the Illumina TruSeq DNA library protocol and sequenced on the Illumina MiSeq (300 bp paired-end). Sequence data (available through the NCBI Short Read Archive PRJNA328211) were analyzed using Quantitative Insights Into Microbial Ecology [QIIME; (63)]. After discarding low-quality sequences (ambiguous base calls, sequences <200 bp and homopolymers >6 bp), barcodes, primers, and chimeras were removed . Operational Taxonomic Units were defined and identified with clustering at 97% similarity and RDP classifier (64) and GreenGenes database (65). After removing chloroplasts, mitochondria, unidentified, and unassigned OTUs, OTU tables were normalized using fourth root transformation and standardized by total by sample. A second OTU table was generated by transforming data into presence/absence. Differences between genotypes and different spatial scales in both datasets were evaluated with a permutational multivariate analysis of variance (PERMANOVA, 9,999 permutations) (66) on Bray-Curtis and Sorensen distances. Core microbiome (100%) was identified for each genotype using QIIME script ‘compute_core_microbiome.py’, and diagrams were generated using Venn diagram software (67).

##### 1.7 Morphological and physiological characterization

Morphological and physiological characterization was undertaken for a subset of samples from the Western Coral Sea. Firstly, gross morphological appearance was visually assessed for colonies with *in situ* photographs of genotyped colonies (n = 157). Bleach-dried skeletons were examined under a stereomicroscope to assess the presence of discriminating, qualitative skeletal characteristics (n = 36; Table S9). Quantitative measurements undertaken for five skeletal characters were measured for a subset of skeletons (n = 54-89): septa per 5 mm, diameter secondary corallite, ridge height, valley width between ridge bottoms, valley width between ridge tops [characters adapted from (68, 69)]. Measurements were conducted in triplicate, except for the diameter of secondary corallite which was dependent on the number of secondary corallites present. Additionally, a small number of representative fragments was then visually assessed using scanning electron microscopy (SEM) to screen for obvious discriminating characters (n = 15).

Physiological characterization was undertaken for a subset of *P. speciosa* samples (n = 73) that were collected from 10, 20, 40 and 60 m depth at three different Osprey sites, and were snap-frozen in liquid nitrogen and subsequently stored at -20 ºC. Coral tissue was removed using an air-brush and 10 mL filtered phosphate buffer. Following centrifugation of the homogenate, the supernatant was frozen (-80 °C) for host protein analysis. The pellet was resuspended in 3 mL filtered phosphate buffer and evenly separated into three aliquots stored at -80 °C until further processing for symbiont cell count and pigment quantification with high-performance liquid chromatography (HPLC), keeping the third sample as a back-up. Symbiodiniaceae density (per $cm^2$) was determined with a Neubauer Improved Bright-Line haemocytometer and a Olympus BX43 light microscope with 6 replicate counts of diluted samples (70, 71). The Symbiodiniaceae counts were normalized against the surface area of the sample skeleton, which was determined by weighing the coral skeleton three times before and after wax dipping. A calibration curve of objects with known surface area was generated to calculate the surface area using the below equation. The Whitaker and Granum (72) methodology was used to obtain the water-soluble protein content (mg $cm^{-2}$) from the host tissue suspension from the air-brushing procedure and standardised to the surface area ($cm^2$). The total lipid concentration (mg $cm^{-2}$) was determined using a modified method of Folch et al. (73) as described by Dunn et al. (74). Frozen coral fragments of 2-3 $cm^2$ were used and resulting lipid content was standardised to the surface area ($cm^2$) of each fragment. Pigment concentration in Symbiodiniaceae cells were determined by high-performance liquid chromatography (HPLC). Pigment concentrations (chlorophyll α, chlorophyll c2, peridin, fucoxanthin, diatoxanthin (Dtx), and diadinoxanthin (Ddx)) were measured with a HPLC using the method as described by Zapata et al. (75) and Dove et al. (76) and normalised to surface area ($cm^2$), Symbiodiniaceae cell count (pg $cell^{-1}$), and chlorophyll α (Chl $\alpha^{-1}$). Lastly, the total xanthophyll pool per chlorophyll α ((Dtx + Ddx) Chl $\alpha^{-1}$) was calculated.

Statistical differences in quantitative morphological and physiological traits were assessed collectively using a permutational multivariate analysis of variance [PERMANOVA (66)], and individually with one-way ANOVA. Three multivariate matrices were generated to evaluate host and symbiont characteristics collectively: morphological traits (septa (per 5mm), width between ridge tops, valley width, ridge height), host (protein and lipids (mg $cm^{-2}$)) and symbiont physiological traits (Chlorophyll α and $c_2$ (Chl $\alpha^{-1}$), Chlorophyll α (pg $cell^{-1}$), Peridin and

Fucoxanthin (Chl $\alpha^{-1}$), Xanthophyll pool (Chl $\alpha^{-1}$) and Symbiodiniaceae ($cm^{-2}$)). Collinearity was evaluated in normalized matrices before the analyses. Differences between lineages and habitats were evaluated on Euclidean distance matrices using Type III sum of squares and 9,999 permutations. The factor "site" was not considered because adequate replication lacked to test it comprehensively (samples for physiology were collected prior to discovering the cryptic diversity), however the three sites were located within 10km of each other. A principal component analysis was performed on the multivariate matrices using the "ggfortify" package in R, and only considering samples with no missing data. Univariate analyses were carried to compare lineages within each depth/habitat (considering those with $n > 3$) with one-way ANOVA (type II) using the package 'car' (77) in R. Non-normal and heteroscedastic data was transformed, unless assumptions were still not met (protein data). Here the ANOVA analysis was still used as it is robust against these violations (78). Significant results were further examined with a Tukey's test (*p*-values were adjusted using single-step method  correction for multiple comparisons).

##### 1.8 Reproductive characterization

Reproductive behavior was assessed at the Orpheus Island Research Station, where 54 large colony fragments were collected from two nearby reef sites between 4-6 November, 2017 (Table S17). Coral colonies were kept in a raceway with flow-through, filtered seawater, constant air bubbling, and a shading canvas to mimic a shaded environment. Colonies were genotyped as described above in the CAPS genotyping section (with a subset of colonies genotyped in the field, and all colonies eventually genotyped in the lab). Between 7-13 November, 2017 colonies were isolated into individual containers and monitored for spawning from 45 minutes before to 2 hours after sunset. Given the predicted split-spawning, 35 of the originally collected coral colonies were kept in a single raceway until the December spawning. Between 7-12 December, 2017, colonies were again isolated into individual containers and monitored for spawning from 30-45 minutes before to 1.5 hours after sunset. Spawning was defined as the vigorous release of gametes (no "dribbling" of either sperm or eggs was observed). All colonies were returned to the reef after spawning observations as per permitting requirements.

#### 2. Supplementary tables

**Table S1. Specimen collection details.** Number of genotyped *Pachyseris speciosa* samples by sampling location (using either nextRAD or CAPS); numbers in brackets refer to *Pachyseris rugosa* samples. Region abbreviations: GBR = Great Barrier Reef (Australia), WCS = Western Coral Sea (Australia), PNG = Kimbe Bay (Papua New Guinea), OKI = Okinawa Islands (Japan), ISR = Eilat (Israel). Habitat abbreviations: B (back-reef, 10 m ± 3), S (reef-slope 10 m ± 3), M (reef-slope 20 m ± 2), D (reef-slope 40 m ± 3), and X (reef-slope 60-85 m). Coordinates are denoted in decimal degrees.

| Region* | Reef | Code | n | Habitats | Latitude | Longitude |
|---|---|---|---|---|---|---|
| GBR | Saunders | GS | 61 (2) | B | S11.44936 | E144.06802 |
| | | | | D | S11.47568 | E144.08582 |
| | Great Detached | GG | 55 (1) | B | S11.70957 | E144.06284 |
| | | | | S,D | S11.70480 | E144.06876 |
| | Mantis | GI | 15 | M,D | S12.21058 † | E143.93278† |
| | Tijou | GT | 143 (1) | S,M,D | S13.06435 | E143.95081 |
| | Tydeman | GY | 74 | S,D | S13.96863 | E144.53009 |
| | Day | GD | 81 (2) | B | S14.51891 | E145.53120 |
| | | | | S,D | S14.47070 | E145.52687 |
| | Ribbon 10 | GR | 60 | B | S14.66926 | E145.65984 |
| | | | | D | S14.67590 | E145.67401 |
| | Ribbon 3 | GB | 19 | S | S15.52752 | 145.779567 |
| | Ruby | GU | 41 (2) | B | S15.74216 | E145.77838 |
| | | | | D | S15.73106 | E145.80539 |
| | Agincourt | GA | 70 (1) | B | S15.99066 | E145.82732 |
| | | | | D | S15.98038 | E145.84122 |
| | Myrmidon | GM | 34 | B | S18.26834 | E147.38144 |
| | | | | D | S18.26567 | E147.40157 |
| | Orpheus Island | GO | 54 | B | S18.66258 | E146.49983 |
| | | | | B | S18.60116 | E146.48841 |
| | | | | B | S18.58131 | E146.49707 |
| | Orpheus – Genome Sample | HS | 1 | B | S18.60800 | E146.48900 |
| WCS | Osprey – Dutch Towers | CO | 100 | S,M,D,X | S13.82032 | E146.56122 |
| | Osprey – Bigeye ledge | CV | 101 | S,M,D,X | S13.85846 | E146.56196 |
| | Osprey – Halfway wall | CY | 177 | S,M,D,X | S13.88925 | E146.55503 |
| | Osprey - Miscellaneous | CZ | 51 | S,D | (between other Osprey sites) | |
| | Bougainville | CB | 71 | S,D,X | S15.49528 | E147.08683 |
| | Holmes | CH | 62 | S,D,X | S16.48211 | E147.87111 |
| | Flinders | CF | 54 | S,D,X | S17.69486 | E148.34336 |
| PNG | Garbuna | PG | 68 | S,D | S5.412892 | E150.09120 |
| | Vanessa | PV | 74 | S,D | S5.425616 | E150.09773 |
| OKI | Kumejima Island | OK | 13 | S,D | N26.31200 | E126.76277 |
| | Sesoko Island | OS | 21 | S,D | N26.62391 | E127.86357 |
| ISR | Dekel Beach | RD | 10 | D | N29.538894 | E34.945777 |
| | IUI Research Station | RU | 10 | D | N29.501719 | E34.917677 |

* GBRMPA permits: G12/35281.1, G14/36802.1 (genome sample), and G14/37294.1. Department of the Environment permits: AU-COM2012-151 AU-COM2013-226, and AU-COM2016-308. Okinawa Prefectural Government permit No. 25-18, issued in 2013. Israel Nature and Parks Authority permit: 40598. † General reef location only, exact coordinates not available.

#### Table S2. Overview of DNA and RNA libraries

| Sample | Genomic DNA | | RNA |
|---|---|---|---|
| Sequencing platform | PacBio SMRT Cell | Illumina HiSeq 2500 | Illumina HiSeq 2500 |
| Number of Reads ( x $10^6$) | 8.03 | 203.6 | 238.7 |
| Read Length | 1 - 23 Kb* | 2 x 250 bp | 2 x 100 bp |
| Insert size (bp) | | 450 | |
| Total Bases ( x $10^9$) | 84 | 101.8 | 47.8 |
| Genome coverage† | 94.8 | 91.9 | |

* = 5-95% range. The longest reads extend to 50 Kb. † = based on estimated genome size of 886.1 Mb

#### Table S3. Estimation of genome size and heterozygosity rate

| Program | k-mer size | Genome size (Mb) | heterozygosity rate (%) |
|---|---|---|---|
| sga.preqc | 31 | 886.1 | |
| GenomeScope | 31 | 749.6 | 1.4 |

#### Table S4. Comparison of genome assembly statistics with other robust coral genomes

| | | *Pachyseris speciosa* (this study) | *Fungia fungites* (13) | *Goniastrea aspera* (13) | *Stylophora pistillata* (79) | *Pocillopora damicornis* (80) | *Orbicella faveolata* (81) |
|---|---|---|---|---|---|---|---|
| Genome | Estimated | 886 | 700 | 860 | 434 | 349 | |
| size (Mb) | Assembled | 984 | 606 | 764 | 400 | 234 | 485 |
| No. contigs / scaffolds | | 2,368 | 7,424 | 5,396 | 5,688 | 4,393 | 1,932 |
| Contig / | N50 | 766 | 323 | 518 | 457 | 326 | 1162 |
| Scaffold | Median | 235 | 13 | 26 | 4 | 2 | 15 |
| size (Kb) | Mean | 415 | 81 | 141 | 70 | 53 | 251 |
| | Largest | 4,615 | 1,804 | 2,896 | 2,969 | 2,168 | 4,771 |
| Gap (Ns)% | | 0 | 13.07 | 10.67 | 10.54 | 3.67 | 26.68 |
| GC% | | 39.56 | 38.41 | 39.29 | 38.54 | 37.82 | 38.99 |

#### Table S5. Overview of de novo gene model statistics

| | | *Pachyseris speciosa* (this study) | *Fungia fungites* (13) | *Goniastrea aspera* (13) | *Stylophora pistillata* (79) | *Pocillopora damicornis* (80) | *Orbicella faveolata* (81) |
|---|---|---|---|---|---|---|---|
| No. Genes | | 39,160 | 38,209 | 35,901 | 25,769 | 26,077 | 25,916 |
| Genic Size (Mb) | | 330 | 245 | 283 | 215 | 146 | 245 |
| % genic | | 33.54 | 40.43 | 37.04 | 53.75 | 62.39 | 50.52 |
| Complete CDS | | 32,194 | 30,578 | 24,932 | 23,060 | 21,366 | 24,551 |
| % completeCDS | | 82.21 | 80.03 | 69.45 | 89.49 | 81.93 | 94.73 |
| Gene | mean | 8,608 | 6,555 | 8,073 | 8,412 | 5,860 | 9,780 |
| (bp) | max | 149,029 | 150,466 | 162,044 | 249,773 | 120,740 | 457,682 |
| mRNA | mean | 1,848 | 1,474 | 1,443 | 2,080 | 1,756 | 2,474 |
| (bp) | max | 40,985 | 41,480 | 43,559 | 48,748 | 64,910 | 46,086 |
| Mean No. exons* | | 6.1 | 5.9 | 6.0 | 7.9 | 7.2 | 6.8 |
| Exon | mean | 303 | 250 | 239 | 264 | 245 | 365 |
| (bp) | max | 15,613 | 17,996 | 14,019 | 20,828 | 15,432 | 23,869 |
| Intron | mean | 1,325 | 1,040 | 1,313 | 919 | 665 | 1,213 |
| (bp) | max | 51,587 | 26,803 | 34,467 | 243,902 | 19,173 | 97,774 |

* = per mRNA

#### Table S6. Functional annotation of robust coral genomes

| | | *Pachyseris speciosa* (this study) | *Fungia fungites* (13) | *Goniastrea aspera* (13) | *Stylophora pistillata* (79) | *Pocillopora damicornis* (80) | *Orbicella faveolata* (81) |
|---|---|---|---|---|---|---|---|
| No. Annotated genes | | 39,160 | 38,209 | 35,901 | 25,769 | 26,077 | 25,916 |
| Swiss-Prot | No. Genes | 24770 | 24760 | 21924 | 17669 | 15628 | 19910 |
| | % | 63.25 | 64.8 | 61.07 | 68.57 | 59.93 | 76.83 |
| PFAM-A | No. Domains | 5041 | 5396 | 5089 | 5376 | 5081 | 5077 |
| | No. Genes | 26373 | 25446 | 22882 | 18778 | 16163 | 19896 |
| | % | 67.35 | 66.60 | 63.74 | 72.87 | 61.98 | 76.77 |

#### Table S7. Genome assembly completeness as benchmarked through universal single-copy orthologs

| | Genome assembly BUSCO (n=978) | | | Predicted transcripts BUSCO (n=978) | | |
|---|---|---|---|---|---|---|
| | % of full | % of partially | % of missing | % of full | % of partially | % of missing |
| *Pachyseris speciosa* (this study) | 81.3 | 1.5 | 17.2 | 90.5 | 5.1 | 4.4 |
| *Fungia fungites* (13) | 86.8 | 4.3 | 8.9 | 90.5 | 6.6 | 2.9 |
| *Goniastrea aspera* (13) | 88.9 | 2.8 | 8.3 | 88.8 | 6.4 | 4.8 |
| *Stylophora pistillata* (79) | 88.2 | 2.9 | 8.9 | 90.4 | 4.2 | 5.4 |
| *Pocillopora damicornis* (80) | 88.3 | 2.9 | 8.8 | 92.4 | 3.4 | 4.2 |
| *Orbicella faveolata* (81) | 85.6 | 4.7 | 9.7 | 90 | 5 | 5 |

#### Table S8. *de novo* annotated repeat content data for robust coral genomes

| | | *Pachyseris speciosa* (this study) | *Fungia fungites* (13) | *Goniastrea aspera* (13) | *Stylophora pistillata* (79) | *Pocillopora damicornis* (80) | *Orbicella faveolata* (81) |
|---|---|---|---|---|---|---|---|
| total length (Mb) | | 984 | 606 | 764 | 400 | 234 | 485 |
| bases masked (Mb) | | 513 | 227 | 341 | 110 | 47 | 112 |
| % bases masked | | 52.19 | 37.54 | 44.69 | 27.5 | 20.26 | 23.14 |
| Inter * | Total | 49.9 | 35.97 | 43.06 | 26.25 | 18.8 | 21.86 |
| | SINEs | 2.22 | 1.42 | 1.77 | 2.12 | 1.08 | 1.07 |
| | LINEs | 8.57 | 2.01 | 3.37 | 4.8 | 2.03 | 1.08 |
| | LTR elements | 2.04 | 0.83 | 1.43 | 0.91 | 0.3 | 0.64 |
| | DNA elements | 12.11 | 8.62 | 9.21 | 4.31 | 2.41 | 5.46 |
| | Unclassified | 25.03 | 23.14 | 27.35 | 14.22 | 13.39 | 13.67 |
| Other ** | Small RNA | 0.3 | 0.01 | 0.08 | 0.04 | 0.19 | 0.17 |
| | Satellites | 0.16 | 0.1 | 0.38 | 0.02 | 0.04 | 0.08 |
| | Simple repeats | 2.02 | 1.33 | 1.16 | 1 | 0.87 | 0.93 |
| | Low complexity | 0.12 | 0.13 | 0.08 | 0.14 | 0.16 | 0.1 |

* = Interspersed repeats (%). ** = Other repeats (%)

**Table S9. Qualitative morphological assessment of *P. speciosa* specimens.** For the "red" lineage a total of 8 skeletons were assessed (5 from 10 m, 3 from 40m), for the "blue" a total of 13 skeletons (5 from 10 m, 5 from 40m, and 3 from 60 m), and for the "green" a total of 15 skeletons (5 from 10 m, 5 from 40m, and 5 from 60 m). Traits 8-11 are adapted from (69).

| Morphological character | Character states | "red" | | | | "green" | | | | "blue" | | | |
|---|---|---|---|---|---|---|---|---|---|---|---|---|---|
| | | a | b | c | d | a | b | c | d | a | b | c | d |
| 1. Unifacial corallum † | a: yes<br>b: no | 15 | 0 | | | 15 | 0 | | | 8 | 0 | | |
| 2. Concentric carinae | a: yes<br>b: mostly<br>c: no | 6 | 2 | 0 | | 13 | 0 | 0 | | 10 | 5 | 0 | |
| 3. Continuous carinae | a: mostly continuous<br>b: few short carinae<br>c: many short carinae | 6 | 0 | 2 | | 11 | 2 | 0 | | 6 | 4 | 5 | |
| 4. Carinae vertical development | a: straight<br>b: mostly straight<br>c: mostly wavy<br>d: wavy | 6 | 1 | 0 | 1 | 8 | 2 | 0 | 3 | 4 | 6 | 1 | 4 |
| 5. Secondary carinae present | a: yes<br>b: few (≤ 3)<br>c: no | 3 | 4 | 1 | | 7 | 6 | | | 10 | 5 | | |
| 6. Secondary calices between secondary carinae | a: yes<br>b: no | 5 | 3 | | | 8 | 5 | | | 14 | 1 | | |
| 7. Secondary calices on sides of carinae | a: yes<br>b: no | 1 | 7 | | | 4 | 9 | | | 5 | 10 | | |
| 8. Columellae mostly fused | a: yes<br>b: no | 5 | 3 | | | 13 | 0 | | | 6 | 9 | | |
| 9. Spatula-shaped processes † | a: yes<br>b: no | 8 | 0 | | | 13 | 0 | | | 15 | 0 | | |
| 10. Zig-zag septo-costae † | a: yes<br>b: no | 8 | 0 | | | 13 | 0 | | | 15 | 0 | | |
| 11. Alternating / equal septo-costae | a: mostly alternating<br>b: mostly equal | 2 | 6 | | | 1 | 12 | | | 4 | 11 | | |

† = No variability in state was observed.

**Table S10. Whole-genome sequencing coverage estimates.** Values are estimates based on scaffold Sc0000001. Samples used for MSMC analysis are indicated with an asterisk (*).

| Sample | Location | Coverage | Lineage | MSMC |
|---|---|---|---|---|
| DC1104 | GBR - Myrmidon BackReef | 4.45 | "green" | |
| DC1107 | GBR - Myrmidon BackReef | 23.1 | "blue" | * |
| DC1108 | GBR - Myrmidon BackReef | 15.2 | "blue" | * |
| DC1105 | GBR - Myrmidon BackReef | 21.1 | "red" | * |
| DC1109 | GBR - Myrmidon BackReef | 28 | "red" | * |
| DC7967 | GBR - GreatDet BackReef | 24.2 | "green" | * |
| DC7969 | GBR - GreatDet BackReef | 22.3 | "green" | * |
| DC7962 | GBR - GreatDet BackReef | 27.8 | "blue" | * |
| DC7968 | GBR - GreatDet BackReef | 26.6 | "blue" | * |
| DC7955 | GBR - GreatDet BackReef | 26.6 | "red" | * |
| DC7957 | GBR - GreatDet BackReef | 28.9 | "red" | * |
| DC7958 | GBR - GreatDet BackReef | 8.6 | "red" | |
| DC8218 | GBR - Ribbon10 BackReef | 6.1 | "green" | |
| DC8230 | GBR - Ribbon10 BackReef | 6.5 | "green" | |
| DC8235 | GBR - Ribbon10 BackReef | 6.5 | "green" | |
| DC8220 | GBR - Ribbon10 BackReef | 5 | "blue" | |
| DC8229 | GBR - Ribbon10 BackReef | 5.6 | "blue" | |
| DC8222 | GBR - Ribbon10 BackReef | 4 | "red" | |
| DC8223 | GBR - Ribbon10 BackReef | 4.4 | "red" | |
| DC8238 | GBR - Ribbon10 BackReef | 4.6 | "red" | |

**Table S11. Primer sequences and enzyme combinations for the CAPS assay**

| Primer name | Primer sequences | Enzyme | Digest: no | Digest: yes |
|---|---|---|---|---|
| Pspe-Green-CfoI/HhaI | F: ACCTGGTGACCTTTGCCATA<br>R: TCTGTCAGTAGAGGGAGGGG | CfoI/HhaI | not "green" | "green" |
| Pspe-Blue-HaeIII | F: CCGTTTCTTCGTCAGCCATT<br>R: CACATCGCTCTTCTTCCGTT | HaeIII | not "blue" | "blue" |
| Pspe-Red-Taqα1 | F: TAATCGCACTGCTAGGGACG<br>R: CTTGGTCTGTTTGTAGCCGT | Taqα1 | "red" | not "red |

**Table S12. Permutational analysis of variance (ADONIS/PERMANOVA) of relative abundances of the three *P. speciosa* lineages across habitats.** Included regions were the Great Barrier Reef (GBR), Western Coral Sea (WCS) and Papua New Guinea (PNG). Test based on Bray-curtis distances with significant differences indicated in bold. For pairwise comparisons, *p*-values were adjusted using Bonferroni correction for multiple comparisons.

| | | Df | F | $R^2$ | *p*-adjusted |
|---|---|---|---|---|---|
| overall | habitat | 1 | 12.5268 | 0.25008 | **0.0009766** |
| | region | 2 | 4.2157 | 0.16832 | **0.0195312** |
| | habitat : region | 2 | 3.066 | 0.12244 | **0.0351562** |
| GBR | habitat | 3 | 5.5909 | 0.46887 | **0.001302** |
| | 40m vs Back | 1 | 9.623429 | 0.3908241 | **0.00400000** |
| | 40m vs 10m | 1 | 4.190730 | 0.2758742 | **0.03500000** |
| | 40m vs 20m | 1 | 3.280852 | 0.2671518 | 0.11000000 |
| | Back vs 10m | 1 | 2.472676 | 0.1982474 | 0.12300000 |
| | Back vs 20m | 1 | 4.824275 | 0.3761831 | **0.03500000** |
| | 10m vs 20m | 1 | 10.364080 | 0.7215276 | 0.06666667 |
| WCS | habitat | 3 | 6.9127 | 0.59699 | **0.0001** |
| | 10m vs 40m | 1 | 11.751378 | 0.5402590 | **0.006** |
| | 10m vs 60m+ | 1 | 9.405625 | 0.5733171 | **0.016** |
| | 10m vs 20m | 1 | 3.559910 | 0.3371155 | 0.083 |
| | 40m vs 60m+ | 1 | 1.364908 | 0.1631707 | 0.309 |
| | 40m vs 20m | 1 | 4.145794 | 0.3719604 | **0.039** |
| | 60m+ vs 20m | 1 | 6.186872 | 0.6073378 | 0.100 |
| PNG | habitat | 1 | 0.20857 | 0.09443 | 1 |

**Table S13. One-factorial ANOVA and pairwise comparisons for relative abundances of the three lineages across habitats.** Included regions were the Great Barrier Reef (GBR) and Western Coral Sea (WCS). The data met the expectations of normal distribution and homoscedasticity (a square root transformation was applied to the relative abundances of the "green" lineage on the GBR). Bold *p*-values indicate significance after single-step correction.

| | "Red" | | | "Blue" | | | "Green" | | |
|---|---|---|---|---|---|---|---|---|---|
| | Df | F | p | Df | F | p | Df | F | p |
| GBR | 3 | 7.917 | **0.001259** | 3 | 3.5965 | **0.03271** | 3 | 4.9463 | **0.01052** |
| WCS | 3 | 11.962 | **0.0003741** | 3 | 4.9228 | **0.01537** | 3 | 2.6333 | 0.09074 |

Pairwise comparisons (Tukey's test)

| | | "Red" | | "Blue" | | "Green" | |
|---|---|---|---|---|---|---|---|
| | Groups | t | *p*-adjusted | t | *p*-adjusted | t | *p*-adjusted |
| GBR | 10m, Back | -2.008 | 0.213 | 2.153 | 0.1670 | 0.087 | 0.9997 |
| | 20m, Back | -1.914 | 0.248 | -1.528 | 0.4322 | 2.796 | **0.0494** |
| | 40m, Back | -4.860 | **<0.001** | 1.497 | 0.4493 | 2.938 | **0.0371** |
| | 20m, 10m | -0.327 | 0.987 | -2.917 | **0.0385** | 2.491 | 0.0900 |
| | 40m, 10m | -1.883 | 0.261 | -0.983 | 0.7541 | 2.287 | 0.1317 |
| | 40m, 20m | 1.085 | 0.695 | 2.476 | 0.0924 | -1.002 | 0.7436 |
| WCS | 20m, 10m | -2.598 | 0.08556 | 0.292 | 0.9909 | | |
| | 40m, 10m | -4.903 | **0.00107** | 2.564 | 0.0913 | | |
| | 60m, 10m | -5.143 | **< 0.001** | 3.347 | **0.0216** | | |
| | 40m, 20m | -1.405 | 0.51351 | 1.802 | 0.3100 | | |
| | 60m, 20m | -2.204 | 0.16813 | 2.646 | 0.0791 | | |
| | 60m, 40m | -1.140 | 0.66919 | 1.253 | 0.6027 | | |

**Table S14. Permutational analysis of variance (PERMANOVA) and pairwise analysis on coral host skeletal traits, coral host physiology and Symbiodiniaceae photophysiological traits (multivariate).** Test based on Euclidean distances on normalized data. Significant differences are indicated in bold. P(perm): *p*-value based on permutations, U. perms: Unique permutations.

| | Coral host skeletal traits | | | Coral host physiology | | | Symbiodiniaceae photophysiological traits | | |
|---|---|---|---|---|---|---|---|---|---|
| Source | Pseudo -F | U. perms | P (perm) | Pseudo -F | U. perms | P (perm) | Pseudo -F | U.perm s | P (perm) |
| Habitat (Ha) | 0.7545 | 9966 | 0.5473 | 3.1304 | 9942 | **0.0076** | 1.4169 | 9925 | 0.1407 |
| Lineages (Li) | 3.6744 | 9932 | **0.0009** | 5.2194 | 9956 | **0.001** | 3.5451 | 9932 | **0.0005** |
| HaxLi | 0.7082 | 9925 | 0.6804 | 1.351 | 9935 | 0.2286 | 1.4517 | 9897 | 0.1037 |

Pairwise - Lineages (Li)

| | Coral host skeletal traits | | | Coral host physiology | | | Symbiodiniaceae photophysiological traits | | |
|---|---|---|---|---|---|---|---|---|---|
| Groups | t | U. perms | P(perm ) | t | U. perms | P(perm ) | t | U. perms | P(perm ) |
| “Green”, “Red” | 2.058 | 9955 | **0.0072** | 2.9382 | 9946 | **0.0001** | 2.4055 | 9955 | **0.0004** |
| “Green”, “Blue” | 0.9927 | 9941 | 0.4295 | 1.5446 | 9955 | 0.103 | 1.9852 | 9941 | **0.0075** |
| “Red”, “Blue” | 2.2762 | 9946 | **0.0018** | 1.3144 | 9947 | 0.1905 | 0.6103 | 9950 | 0.8779 |

Pairwise - Habitats (Ha), coral host physiology

| Groups | t | U. perms | P(perm) |
|---|---|---|---|
| 10m, 40m | 0.93723 | 9948 | 0.4186 |
| 10m, 20m | 0.74763 | 9945 | 0.5821 |
| 10m, 60m | 1.2155 | 9965 | 0.2461 |
| 40m, 20m | 1.9483 | 9955 | 0.0221 |
| 40m, 60m | 2.7635 | 9931 | **0.0074** |
| 20m, 60m | 2.2177 | 9950 | **0.0113** |

**Table S15. One-factorial ANOVA for physiological and morphological parameters between genetic lineages for each depth.** † indicates that data was non-normal but 1-way ANOVA was still used as it is robust against this violation *(Harwell et al. 1992)*. ‡ indicates that data was Log10 transformed to meet assumptions of normal distribution and homoscedasticity. Significance after correction (single-step method) is indicated in bold *p*-values for the main analysis and with asterisks for pairwise comparisons (* = *p*<0.05, ** = *p* <0.01).

| Skeletal traits | Between lineages (10 m) | | | Between lineages (40 m) | | |
|---|---|---|---|---|---|---|
| Septa per 5 mm | F(2)=5.007 | **p=0.01667** | R<G=B | F(2)=3.7373 | p=0.03691 | n.s. |
| Diameter secondary corallite | F(2)=1.5362 | p=0.2436 | | F(2)=1.179 | p=0.3367 | |
| Ridge height | F(2)=0.831 | p=0.4495 | | F(2)=0.0582 | p=0.9436 | |
| Valley width between ridge bottoms | F(2)=0.8559 | p=0.4392 | | F(2)=2.892 | p= 0.07277 | |
| Valley width between ridge tops | F(2)=2.0737 | p=0.1507 | | F(2)=3.3338 | p=0.05082 | |

| Host physiology | Between lineages (10 m) | | | Between lineages (20 m) | |
|---|---|---|---|---|---|
| Proteins | F(1)=10.421 † | **p=0.00417** | G<R | F(2)=0.9813 | p=0.3914 |
| Lipids | F(1)=1.8756 | p=0.1887 | | F(2)=0.7438 | p=0.4874 |

| Host physiology | Between lineages (40 m) | | | Between lineages (60 m) | | |
|---|---|---|---|---|---|---|
| Proteins | F(2)=2.4333 | p=0.1195 | | F(1)=9.231 | **p=0.02881** | B>G |
| Lipids | F(2)=3.2729 | p=0.06434 | | F(1)=1.4023 | p=0.2896 | |

| Symbiont physiology | Between lineages (10 m) | | | Between lineages (20 m) | | |
|---|---|---|---|---|---|---|
| Symbiodiniaceae density | F(1)=6.5189 ‡ | **p=0.02057** | G<R | F(2)=0.4388 ‡ | p=0.6506 | |
| Chlorophyll α (μg $cm^{-2}$) | F(1)=0.2024 | p=0.6585 | | F(2)=0.1994 | p=0.8207 | |
| Chlorophyll α (pg $cell^{-1}$) | F(1)=15 | **p=0.00122** | G>R | F(2)=0.1839 ‡ | p=0.8333 | |
| Total Xanthophyll pool | F(1)=1.1073 | p=0.3074 | | F(2)=7.7445 | **p=0.00302** | G>R ** |
| Fucoxanthin | F(1)=0.2 | p=0.6604 | | F(2)=0.0622 | p=0.9398 | |
| Peridin | F(1)=0.1133 | p=0.7405 | | F(2)=8.9198 | **p=0.00157** | G>B=R ** |
| Chlorophyll $c_2$ | F(1)=14.481 | **p=0.00141** | G<R | F(2)=0.3301 | p=0.7225 | |

| Symbiont physiology | Between lineages (40 m) | | | Between lineages (60 m) | | |
|---|---|---|---|---|---|---|
| Symbiodiniaceae density | F(2)=5.2167 | **p=0.01712** | R>G* | F(1)=1.5893 | p=0.2631 | |
| Chlorophyll α (μg $cm^{-2}$) | F(2)=1.3375 | p=0.2888 | | F(1)=16.403 | **p=0.00982** | G>B |
| Chlorophyll α (pg $cell^{-1}$) | F(2)=0.7316 | p=0.4957 | | F(1)=0.4744 | p=0.5216 | |
| Total Xanthophyll pool | F(2)=6.3886 | **p=0.00852** | G > R=B* | F(1)=2.3084 | p=0.1891 | |
| Fucoxanthin | F(2)=3.4342 | p=0.05589 | | F(1)=0.4094 | p=0.5504 | |
| Peridin | F(2)=1.5686 | p=0.237 | | F(1)=2.561 | p=0.1704 | |
| Chlorophyll $c_2$ | F(2)=0.0756 | p=0.9275 | | F(1)=0.1771 | p=0.6913 | |

**Table S16. Permutational analysis of variance (PERMANOVA) on bacterial community.** Test based on: a) structure: Bray-Curtis distance on fourth root transformed and standardized data, composition: Sorensen distance on presence/absence data, c) richness, diversity, Delta+ and Lambda+: Euclidean distance. Significant differences are indicated in bold. P(perm): *P*-value based on permutations, U. perms: Unique permutations.

| | Structure | | | Composition | | | Richness | | |
|---|---|---|---|---|---|---|---|---|---|
| Source | Pseudo -F | U. perms | P (perm) | Pseudo -F | U. perms | P (perm) | Pseudo -F | U. perms | P (perm) |
| Region (Re) | 1.4015 | 9814 | **0.0038** | 1.5934 | 9820 | **0.0001** | 0.4182 | 9799 | 0.4918 |
| Habitat (Ha) | 0.8770 | 9827 | 0.7898 | 0.8890 | 9845 | 0.7643 | 0.2605 | 9789 | 0.5722 |
| Lineages (Li) | 1.0225 | 9734 | 0.4166 | 1.0085 | 9695 | 0.4713 | 0.0978 | 9954 | 0.9015 |
| RexHa | 1.0095 | 9795 | 0.4679 | 1.0276 | 9796 | 0.4166 | 5.7533 | 9834 | **0.025** |
| RexLi | 1.0274 | 9731 | 0.3917 | 1.0129 | 9714 | 0.4543 | 1.6598 | 9945 | 0.2002 |
| HaxLi | 1.066 | 9735 | 0.2535 | 1.0783 | 9703 | 0.2126 | 0.9160 | 9946 | 0.3714 |
| RexHaxLi | 0.9725 | 9798 | 0.5746 | 0.9639 | 9802 | 0.5996 | 2.6954 | 9838 | 0.1131 |

| | Diversity | | | Delta+ | | | Lambda+ | | |
|---|---|---|---|---|---|---|---|---|---|
| Source | Pseudo -F | U. perms | P (perm) | Pseudo -F | U. perms | P (perm) | Pseudo -F | U. perms | P (perm) |
| Region (Re) | 0.0372 | 9848 | 0.8546 | 1.9357 | 9854 | 0.1691 | 0.6625 | 9836 | 0.4209 |
| Habitat (Ha) | 0.0308 | 9825 | 0.8586 | 0.7479 | 9834 | 0.3918 | 0.8320 | 9843 | 0.3608 |
| Lineages (Li) | 0.3749 | 9938 | 0.6828 | 1.2611 | 9948 | 0.2959 | 1.663 | 9948 | 0.2018 |
| RexHa | 5.5463 | 9850 | **0.0236** | 0.4231 | 9839 | 0.5272 | 1.7815 | 9820 | 0.193 |
| RexLi | 1.2622 | 9941 | 0.3047 | 1.588 | 9943 | 0.2218 | 1.3628 | 9933 | 0.2685 |
| HaxLi | 0.1245 | 9958 | 0.8872 | 0.4851 | 9952 | 0.6169 | 0.5825 | 9943 | 0.5498 |
| RexHaxLi | 1.7416 | 9822 | 0.197 | 0.0608 | 9817 | 0.8038 | 0.0132 | 9850 | 0.91 |

Pairwise - Region * Habitat

| | Richness | | | Diversity | | |
|---|---|---|---|---|---|---|
| Groups | t | U. perms | P(perm) | t | U. perms | P(perm) |
| Shallow - WCS, GBR | 2.2812 | 9854 | **0.0465** | 2.1762 | 9868 | **0.0493** |
| Deep - WCS, GBR | 1.6549 | 9843 | 0.113 | 1.9875 | 9850 | 0.0582 |

**Table S17. Spawning observations from Orpheus Island (Great Barrier Reef)**. Numbers refer to the number of colonies releasing sperm, eggs, or no gametes (none).

| Date | Day | "Blue" lineage | | | "Red" lineage | | | "Green" lineage | | |
|---|---|---|---|---|---|---|---|---|---|---|
| | | Sperm | Eggs | None | Sperm | Eggs | None | Sperm | Eggs | None |
| 7-Nov-17 | 3 | 0 | 0 | 0 | 0 | 0 | 0 | 0 | 0 | 0 |
| 8-Nov-17 | 4 | 0 | 0 | 7 | 0 | 1 | 19 | 0 | 0 | 27 |
| 9-Nov-17 | 5 | 0 | 0 | 7 | 8 | 3 | 9 | 0 | 0 | 27 |
| 10-Nov-17 | 6 | 1 | 0 | 6 | 0 | 0 | 20 | 0 | 0 | 27 |
| 11-Nov-17 | 7 | 3 | 0 | 4 | 0 | 0 | 20 | 0 | 0 | 27 |
| 12-Nov-17 | 8 | 0 | 0 | 7 | 0 | 0 | 20 | 0 | 0 | 27 |
| 13-Nov-17 | 9 | 0 | 0 | 7 | 0 | 0 | 20 | 0 | 0 | 27 |
| 7-Dec-17 | 3 | 0 | 0 | 6 | 1 | 0 | 14 | 0 | 0 | 15 |
| 8-Dec-17 | 4 | 1 | 0 | 5 | 5 | 3 | 7 | 0 | 1 | 14 |
| 9-Dec-17 | 5 | 0 | 0 | 6 | 0 | 3 | 12 | 0 | 0 | 15 |
| 10-Dec-17 | 6 | 0 | 0 | 6 | 3 | 3 | 9 | 0 | 0 | 15 |
| 11-Dec-17 | 7 | 0 | 0 | 6 | 0 | 0 | 15 | 0 | 0 | 15 |
| 12-Dec-17 | 8 | 0 | 0 | 6 | 1 | 0 | 14 | 0 | 0 | 15 |

**Table S18. Spawning time observations for individual colonies**

| Sample | lineage | date | gamete type | time of spawning | time of sunset | relative spawning time |
|---|---|---|---|---|---|---|
| DC9166 | "red" | 8-Nov-17 | eggs | 19.15 | 18.24 | 51 |
| DC9188 | "red" | 9-Nov-17 | sperm | 18.29 | 18.25 | 4 |
| DC9156 | "red" | 9-Nov-17 | sperm | 18.3 | 18.25 | 5 |
| DC9180 | "red" | 9-Nov-17 | sperm | 18.31 | 18.25 | 6 |
| DC9164 | "red" | 9-Nov-17 | sperm | 18.31 | 18.25 | 6 |
| DC9163 | "red" | 9-Nov-17 | sperm | 18.4 | 18.25 | 15 |
| DC9159 | "red" | 9-Nov-17 | eggs | 18.35 | 18.25 | 10 |
| DC9175 | "red" | 9-Nov-17 | sperm | 18.43 | 18.25 | 18 |
| DC9204 | "red" | 9-Nov-17 | sperm | 18.57 | 18.25 | 32 |
| DC9166 | "red" | 9-Nov-17 | eggs | 19.17 | 18.25 | 52 |
| DC9160 | "red" | 9-Nov-17 | eggs | 19.23 | 18.25 | 58 |
| DC9179 | "red" | 9-Nov-17 | sperm | 19.26 | 18.25 | 61 |
| DC9191 | "blue" | 10-Nov-17 | sperm | 18.25 | 18.25 | 0 |
| DC9194 | "blue" | 11-Nov-17 | sperm | 18.39 | 18.26 | 13 |
| DC9191 | "blue" | 11-Nov-17 | sperm | 18.39 | 18.26 | 13 |
| DC9157 | "blue" | 11-Nov-17 | sperm | 18.58 | 18.26 | 32 |

##### 3. Supplementary figures

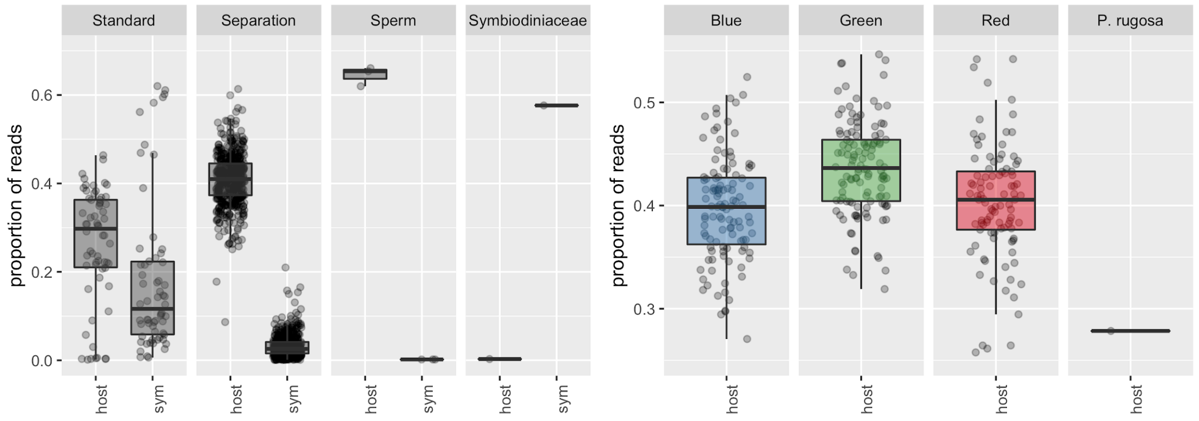

**Figure S1. Proportion of nextRAD reads mapping to genome references.** (left) Proportion of reads mapping to the *Pachyseris speciosa* ("host") and *Cladocopium goreaui* ("sym") genome [29] for the following genomic DNA extraction types: "standard", involving a "separation" step to reduce Symbiodiniaceae contamination, replicate "sperm" samples (as used for the reference genome; from one single colony), and a FACS-isolated Symbiodiniaceae sample. (right) Proportion of reads mapping to the *P. speciosa* genome ("green" lineage) for the main lineages encountered in this study (and three *P. rugosa* outgroup samples).

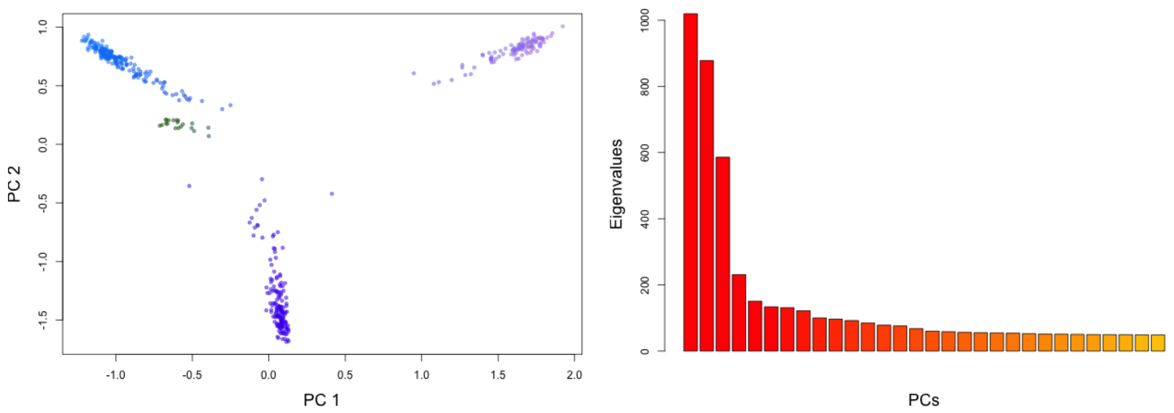

**Figure S2. Principal component analysis (PCA) of *P. speciosa* samples based on nextRAD sequencing.** PCA includes a total of 465 samples (potential clones were removed), with point colors visualizing the contribution of the third principal component (by mapping the first three PCs to RGB color values). Eigenvalues of the PCs are indicated in the bar graph right of the PCA. Jointly, the first three PCs indicate the separation of the overall dataset into four distinct clusters.

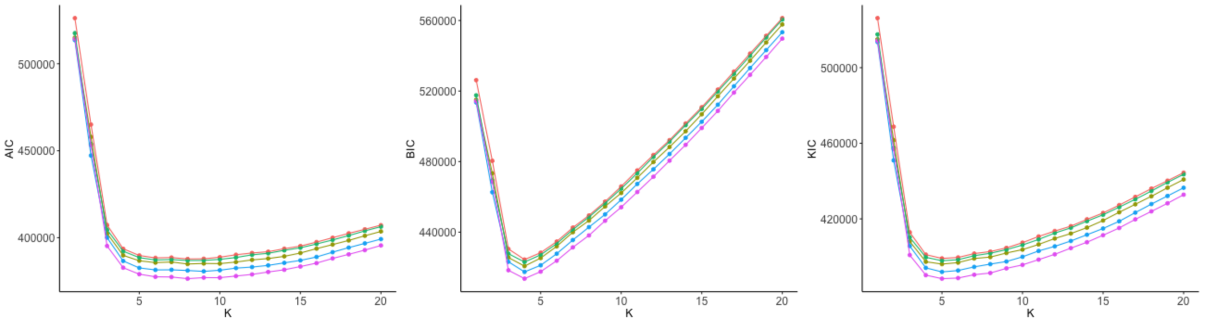

**Figure S3. Goodness-of-fit statistics for increasing numbers of *P. speciosa* clusters in snapclust based on nextRAD sequencing.** The three graphs show the Akaike Information Criterion (AIC), Bayesian Information Criterion (BIC), and Kullback Information Criterion (KIC) for k values ranging from 1 to 20. Jointly they point towards an optimal k of around 4 to 6 clusters.

| | BL2_O | BLU_C | BLU_G | GR2_O | GRN_C | GRN_G | GRN_P | ISR_R | RED_C |
|---|---|---|---|---|---|---|---|---|---|
| BLU_C | 0.0646 | | | | | | | | |
| BLU_G | 0.0642 | 0.0162 | | | | | | | |
| GR2_O | 0.1509 | 0.1116 | 0.1107 | | | | | | |
| GRN_C | 0.1416 | 0.1102 | 0.111 | 0.0967 | | | | | |
| GRN_G | 0.144 | 0.111 | 0.1032 | 0.0881 | 0.0179 | | | | |
| GRN_P | 0.1592 | 0.123 | 0.1261 | 0.1138 | 0.0339 | 0.0225 | | | |
| ISR_R | 0.1839 | 0.1338 | 0.1308 | 0.2225 | 0.1731 | 0.1585 | 0.2136 | | |
| RED_C | 0.1583 | 0.1265 | 0.1314 | 0.1576 | 0.134 | 0.1425 | 0.1476 | 0.2118 | |
| RED_G | 0.1638 | 0.1221 | 0.114 | 0.1574 | 0.134 | 0.1113 | 0.1613 | 0.1912 | 0.013 |

**Figure S4. Genome-wide $F_{ST}$ values for pairwise comparisons of *P. speciosa* populations based on nextRAD sequencing.** Pairwise $F_{ST}$ values (Weir and Cockerham) are calculated for the six lineages ("BLU" = "blue" lineage, "GRN" = "green" lineage, "RED" = "red" lineage, "GR2" = "green" lineage from Okinawa, "BL2" = "blue" lineage from Okinawa, "ISR" = lineage from Israel) and the five different geographic regions in which they occur (C = "Western Coral Sea", G = "Great Barrier Reef", P = "Papua New Guinea", O = "Okinawa", R = "Red Sea). Note that potentially admixed samples (as listed in Figure S5) were excluded from these calculations (as the aim was to quantify differentiation between "purebred" lineages).

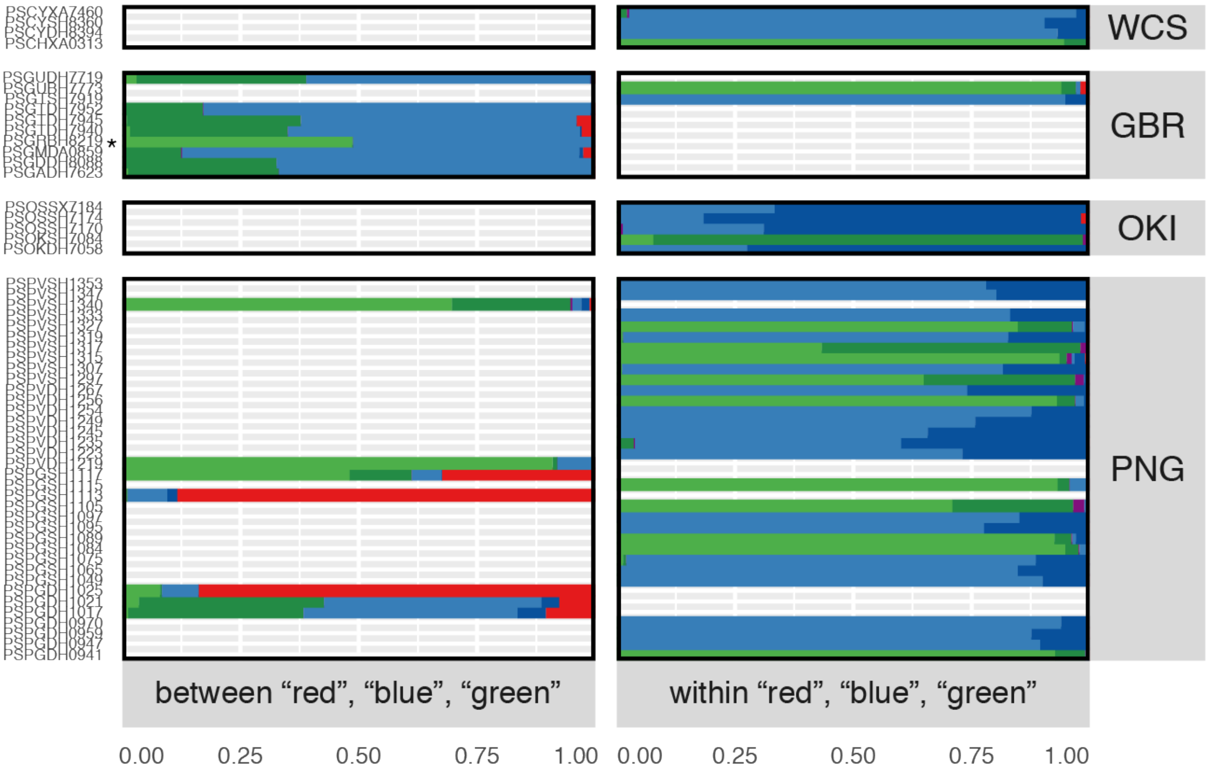

**Figure S5. Potentially admixed samples of *P. speciosa* based on nextRAD sequencing.** Cluster assignments based on STRUCTURE (k = 6) for samples that are potentially admixed (maximum assignment < 0.95 across clusters). The left panel ("between") shows samples that have a mixed assignment between "red", "green" or "blue" clusters, whereas the right panel ("within") shows samples with a mixed assignment within the "green" and "blue" clusters. Note that samples that were fully assigned to the dark-green and dark-blue clusters originated from Okinawa (Figure 2A). The region codes follow those in the manuscript (WCS = Western Coral Sea, GBR = Great Barrier Reef, OKI = Okinawa, and PNG = Papua New Guinea). The seven "admixed" corals observed for Papua New Guinea had assignments to three or more clusters (left panel), likely reflecting clustering artefacts (e.g. due to admixture with an unsampled population/lineage), except for maybe two of the samples (PSPVDH1219 and PSPGSH1113). Seven samples from the Great Barrier Reef showed admixture of the "blue" lineage with the "dark green" cluster present only in Okinawa, which again likely represents a clustering artefact. The F1 hybrid between the "blue" and "green" cluster is indicated with an asterisk.

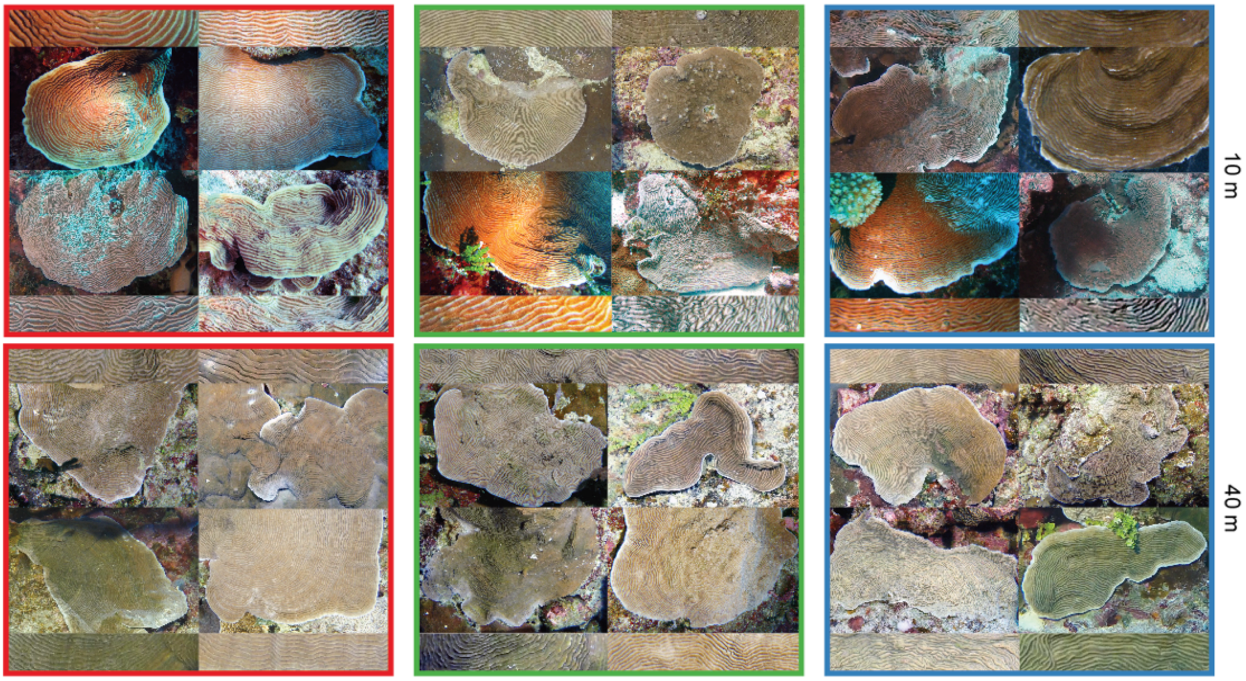

**Figure S6. Morphological variation within the three *P. speciosa* lineages.** Composite shows the gross morphology (and accompanying close-up) of four representative *P. speciosa* colonies from Australia for each of the three lineages ("red", "green", and "blue") from 10 and 40 m depth.

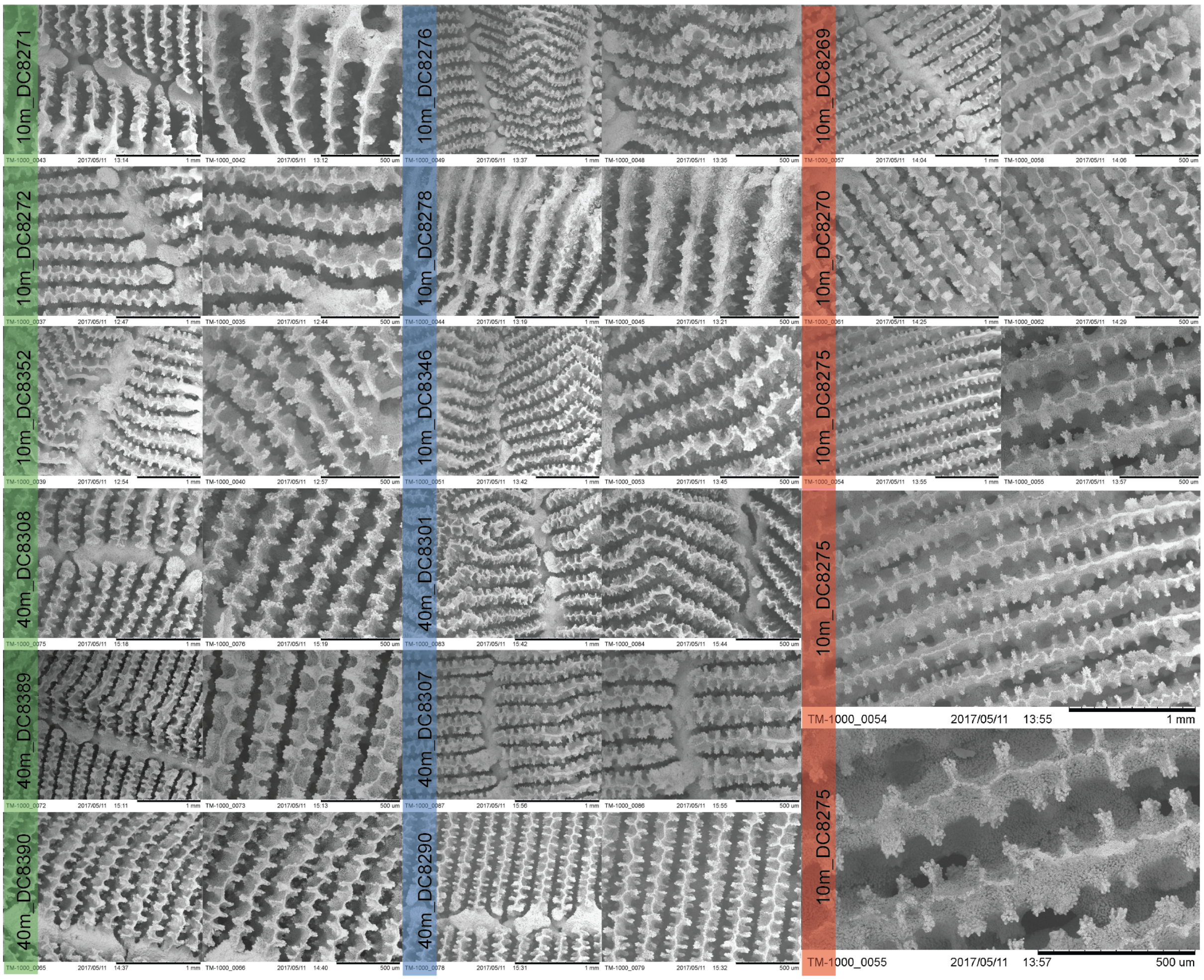

**Figure S7. Scanning electron microscope (SEM) photographs of the three *P. speciosa* lineages.** Composite shows skeletal details (at two different magnifications) for a total of 15 colonies from Australia (6 “green”, 6 “blue” and 3 “red” colonies). Depth, sample number and lineage (color) are indicated for each pair of photographs, and scale is indicated below each photograph.

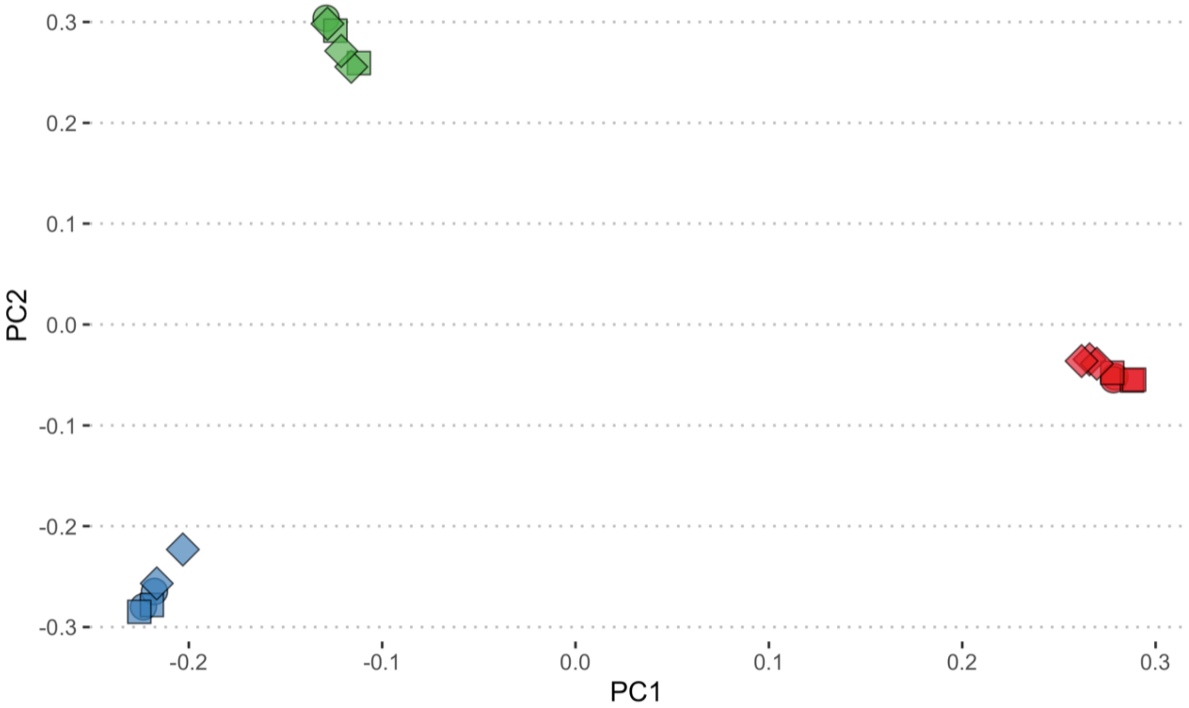

**Figure S8. Principal component analysis (PCA) of *P. speciosa* samples based on whole-genome sequencing.** Genetic structure based on genome-wide genotype likelihoods calculated with PCAngsd. The PCA depicts 20 samples with points coloured according to the three lineages ("red", "green" and "blue"), and shapes corresponding to Great Barrier Reef locations (circle = "Great Detached", triangle = "Myrmidon", and square = "Ribbon Reef 10").

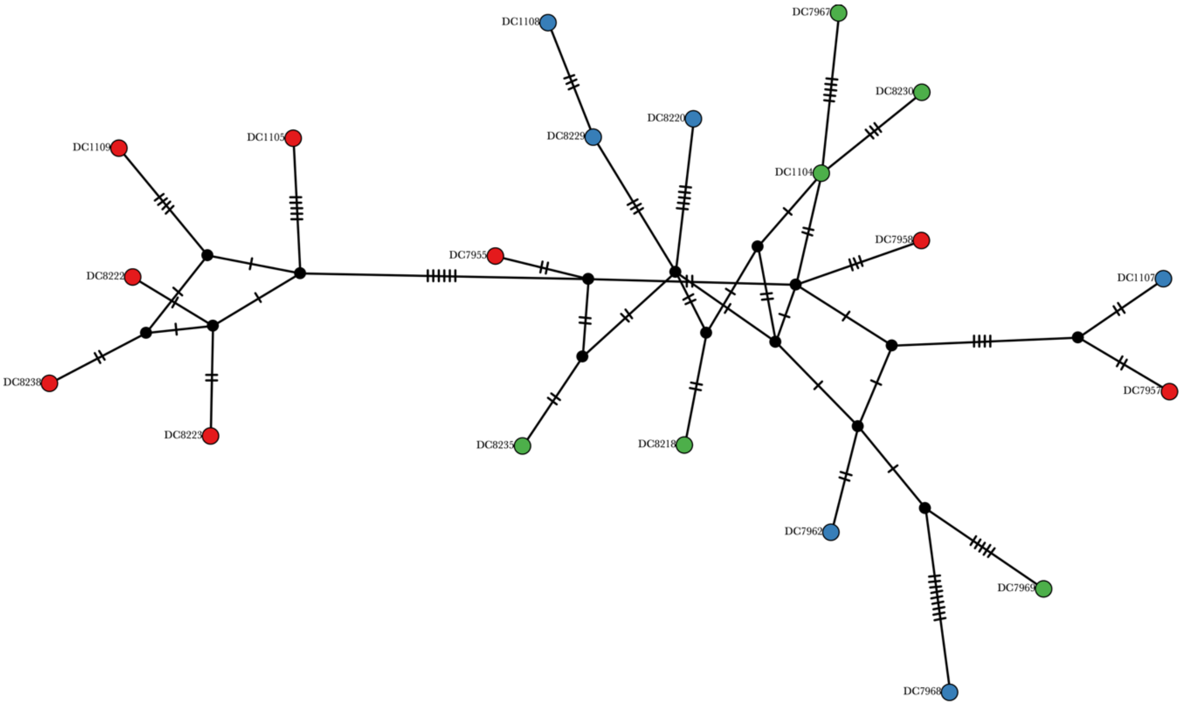

**Figure S9. TCS haplotype network for *P. speciosa* mitochondrial genomes reconstructed from whole-genome sequencing data.** Colours indicate the "red", "green" and "blue" lineages with inferred nodes in black. The network includes a total of 20 samples from the Great Barrier Reef, with all nodes displayed at the same size because no shared haplotypes were present (n = 20). Bars across edges indicate numbers of mutations between nodes.

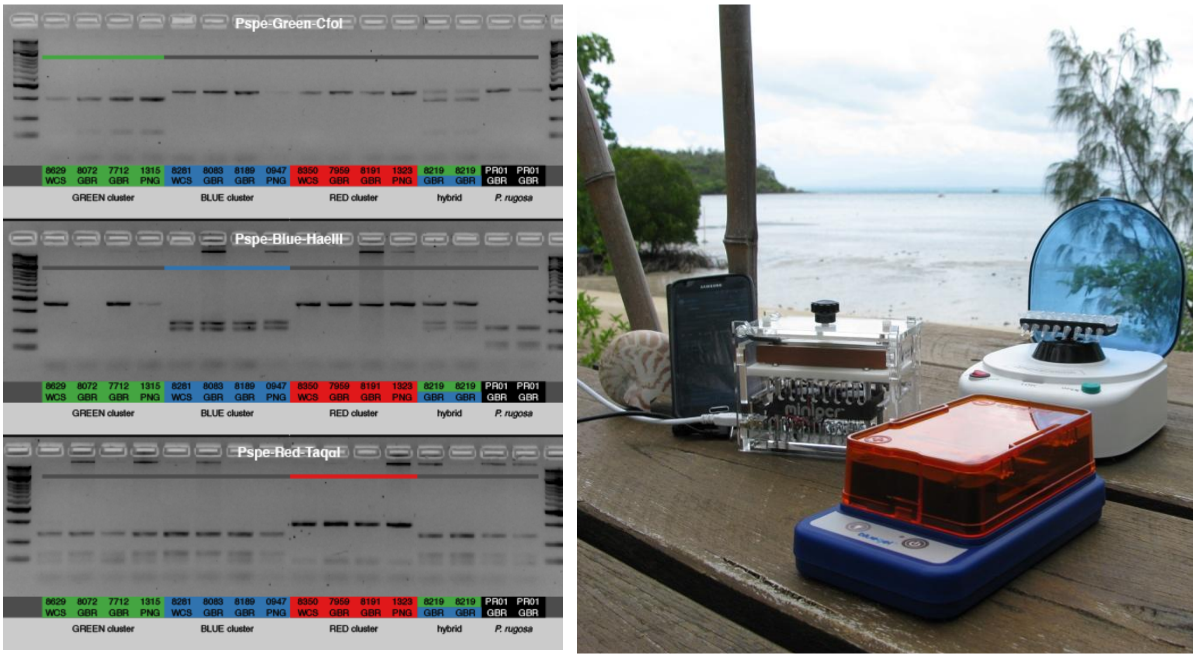

**Figure S10. Cleaved amplified polymorphic sequence (CAPS) used to genotype *P. speciosa* lineages.** (left) Example of diagnostic amplicons for the three different CAPS markers, using the same 15 samples. (right) Field-based genotyping set-up as used for the identification of spawning corals at Orpheus Island on the Great Barrier Reef (Photo by Dagmar Wels).

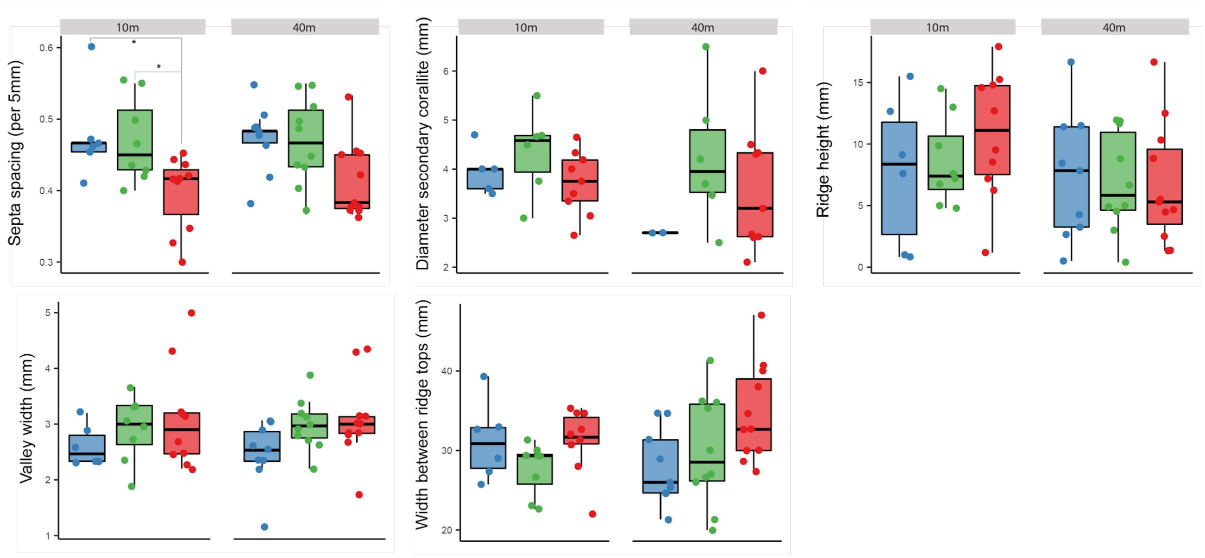

**Figure S11. Comparison of five skeletal traits between the three *P. speciosa* lineages at different depths.** Measurements for the traits were undertaken for at least 54 Australian samples, except for "diameter secondary corallite" which was measured in 37 samples. Gray lines indicate significant differences, where $p < 0.05$ is indicated by *.

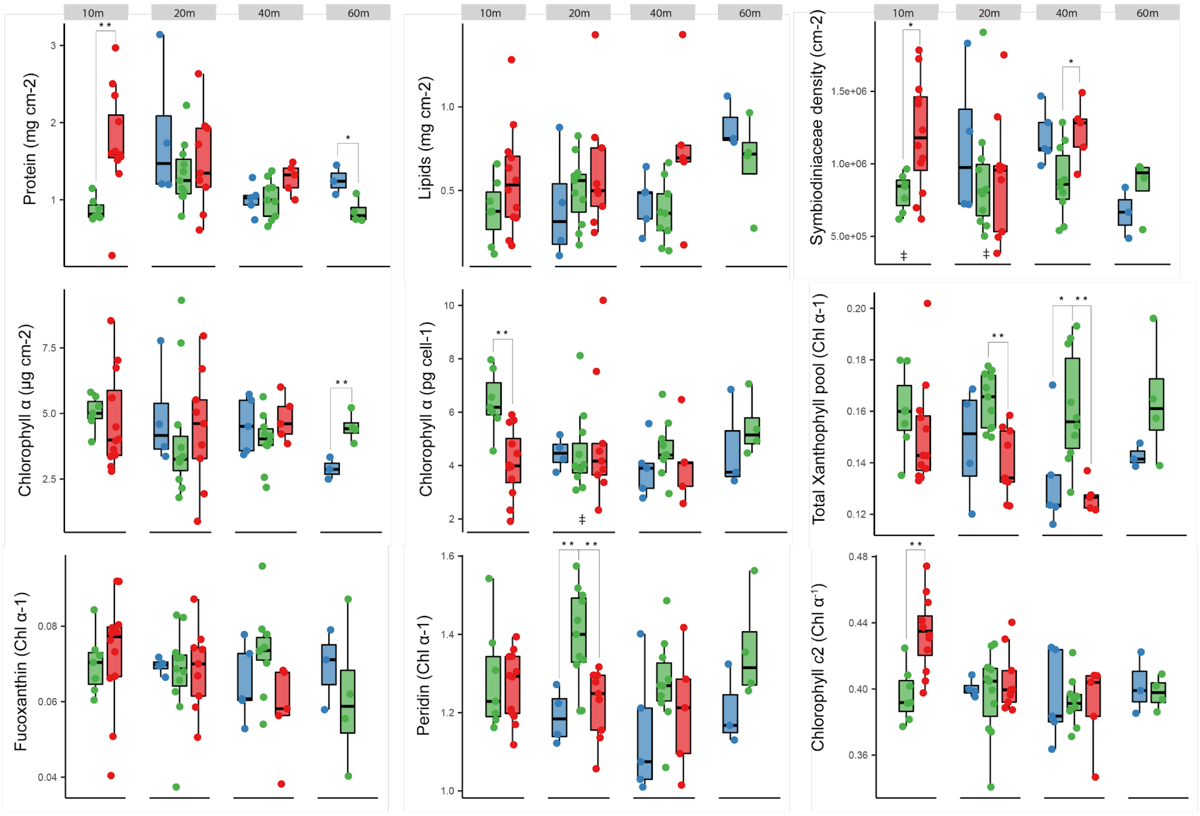

**Figure S12. Comparison of nine physiological traits between the three *P. speciosa* lineages at different depths.** Measurements for each trait were undertaken for 73 samples from Osprey Reef (Western Coral Sea). Gray lines indicate significant differences, where $p < 0.05$ is indicated by *, and $p < 0.01$ is indicated by **.

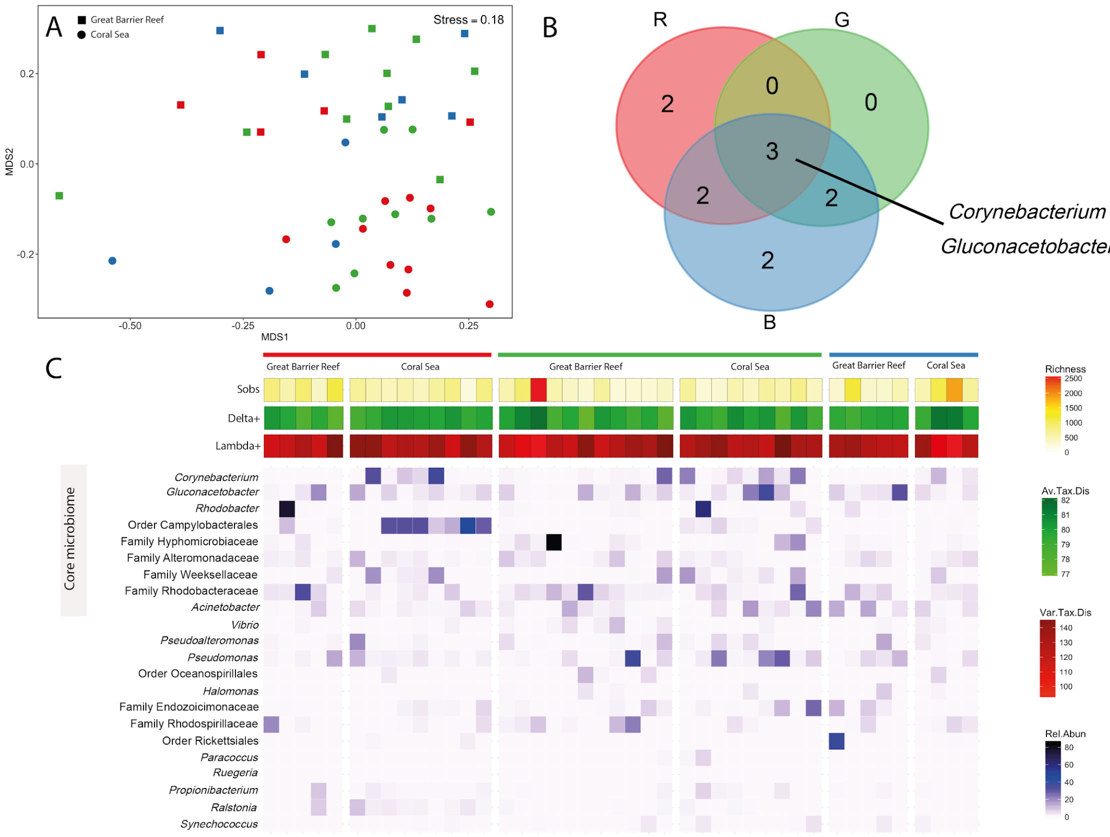

**Figure S13. Characterization of bacterial communities associated with the three *P. speciosa* lineages.** (A) Non-metric multidimensional scaling (NMDS) based on bacterial abundance of 43 samples, showing differences in the structure of coral microbiome occurring between regions rather than between lineages (PERMANOVA, $p < 0.05$). (B) Three bacteria OTUs from the genera *Corynebacterium* and *Gluconacetobacter* were consistently found in the three lineages (intersection of Venn diagram). Lineages "blue" and "red" showed unique persistent OTUs (absent in the other lineages), respectively belonging to genus *Acinetobacter* and family Rhodobacteraceae, and family Weeksellaceae and order Campylobacterales. (C) *P. speciosa* lineages were similar in richness and taxonomic composition (Delta+ and Alpha+); and no differences were identified in the relative abundance of the core microbiome and dominant bacterial OTUs from Alpha- and Gammaproteobacteria classes.

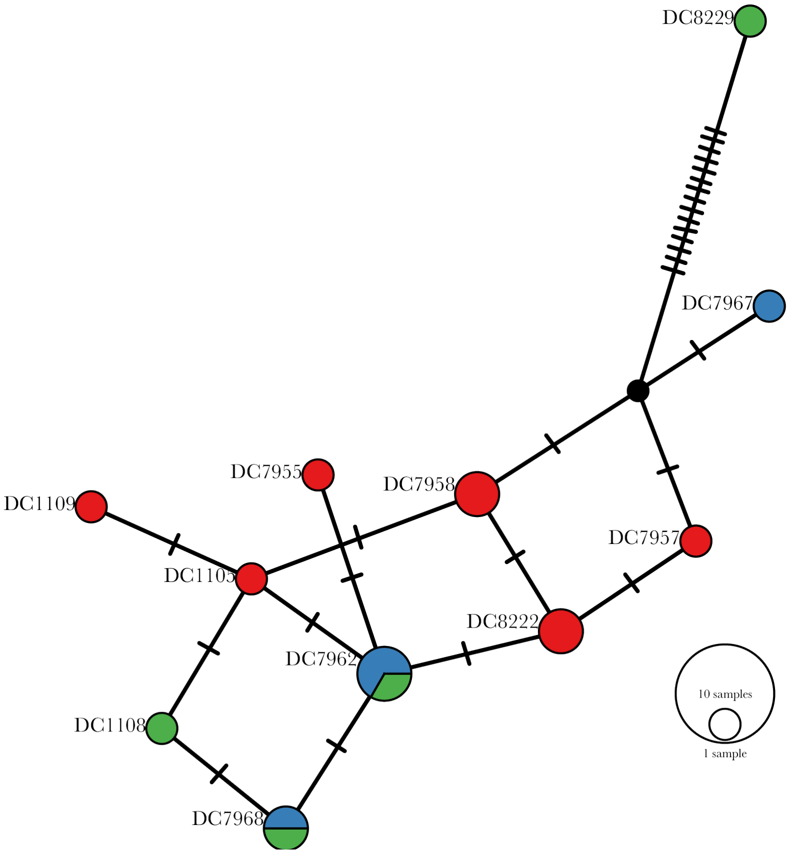

**Figure S14. Haplotype network (TCS) of Symbiodiniaceae mitochondrial genomes associated with the three *P. speciosa* lineages.** Colors correspond to the "red", "green" and "blue" host lineages and cross bars represent the numbers of mutations separating haplotypes. The network includes a total of 16 samples from the Great Barrier Reef, and circle sizes indicate the number of samples with that haplotype.

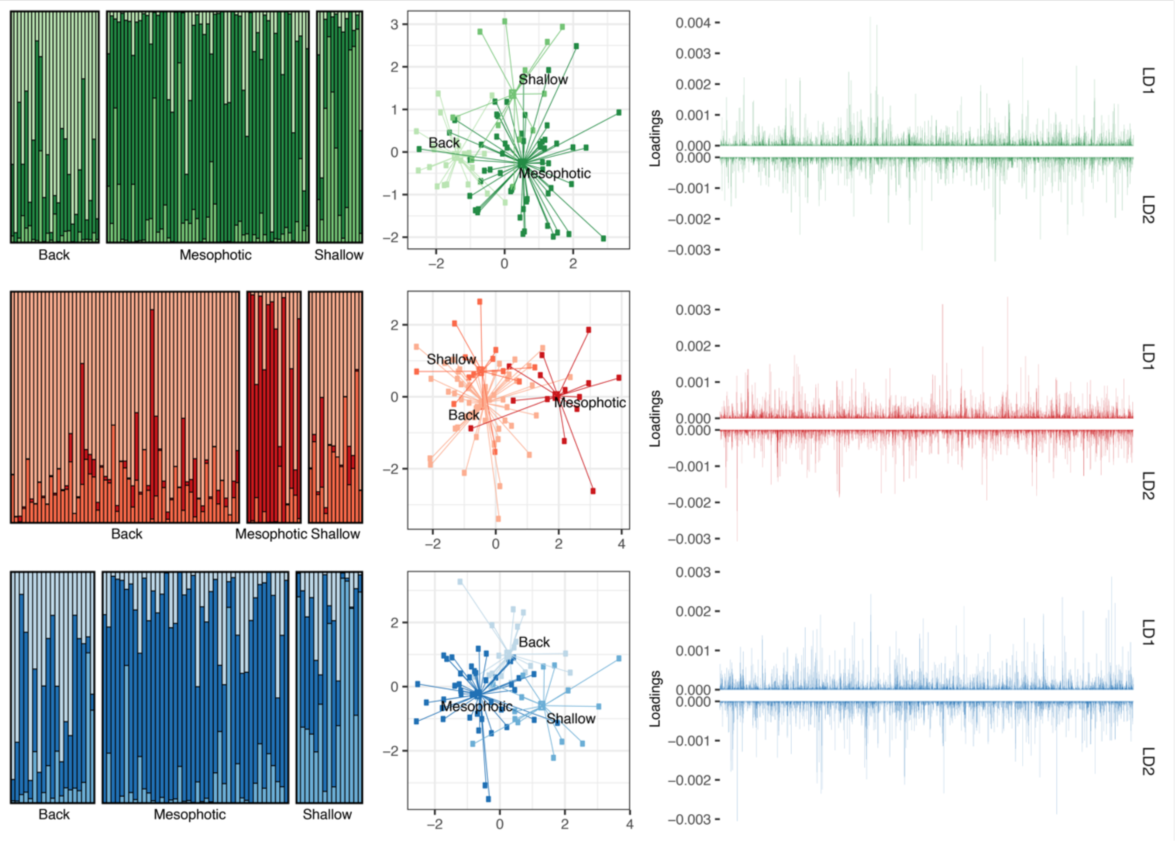

**Figure S15. Genetic structuring across habitats on the Great Barrier Reef (GBR).** (left) DAPC assignment plots for each of the three lineages (from top to bottom: "green", "red" and "blue") using habitat as prior (ignoring locations on the Great Barrier Reef). (middle) DAPC density plot for the Great Barrier Reef using habitat as prior. Plot shows individual colonies connected to respectively region and habitat centroids. (right) DAPC loading plots showing the allele contributions to the first (as positive values) and second (as negative values) discriminant functions when using habitat as prior. Analyses are based on nextRAD data after the removal of outliers.
